# Probing the proteome-wide impact of inhibitors of Leucine-rich Repeat Kinases 1 and 2 on protein-protein interactions and phosphorylation

**DOI:** 10.1101/2025.09.03.674114

**Authors:** Jasmin Jansen, Marcel Wendel, Nicolai D. Raig, Tom Kraus, Katherine J. Surridge, Swetha Mahesula, Nicole Richter-Müller, Sebastian Mathea, Samara L. Reck-Peterson, Stefan Knapp, Florian Stengel

**Affiliations:** Aligning Science Across Parkinson’s (ASAP) Collaborative Research Network, Chevy Chase, MD 20815, USA; Department of Biology, University of Konstanz, Konstanz, Germany; Konstanz Research School of Chemical Biology, University of Konstanz, Germany; Institute of Pharmaceutical Chemistry, Goethe-Universität, Frankfurt 60438, Germany; Structural Genomics Consortium (SGC), Buchmann Institute for Life Sciences, Goethe-Universität, Frankfurt 60438, Germany; Department of Cell and Developmental Biology, School of Biological Sciences, University of California San Diego, La Jolla, CA USA; Department of Cellular and Molecular Medicine, School of Medicine, University of California San Diego, La Jolla, CA USA; Howard Hughes Medical Institute, Chevy Chase, MD 20815, USA

## Abstract

The Leucine-rich repeat kinases 1 and 2 (LRRK1 and 2) are large, multidomain proteins and closely related members of the Roco protein family. They share a high similarity in domain structure and are both phosphorylate members of the Rab GTPase family. However, despite these similarities, there are substantial differences between the two kinases. While mutations to LRRK1 are only implicated in rare cases of osteopetrosis, LRRK2 is associated with multiple diseases, most prominently with familial and sporadic forms of Parkinson’s disease, where pathogenic LRRK2 is associated with an increased kinase activity. While LRRK2 has received major attention from the research community, LRRK1 has been largely understudied. In this work, we employ proximity labelling mass spectrometry in combination with quantitative phosphoproteomics in a model cell line to obtain the cellular interactomes of LRRK1 and LRRK2 and corresponding phosphorylation sites. We then use this dataset to characterize the impact of small molecules targeting both LRRK1 and 2. We identify phosphorylation sites across the proteome that are impacted by these inhibitors and identify novel candidate substrates for LRRK2, including MICALL2. Taken together our data provide a powerful resource for future studies on the cellular role and function of LRRK proteins and their potential use as therapeutic targets.

## Introduction

The Leucine rich-repeat kinases 1 and 2 (LRRK1 and LRRK2) are closely related members within the Roco protein family of GTPases (Marín, 2006; Wauters, Versées and Kortholt, 2019). The two homologs share a similar domain structure with Ankyrin and Leucine-Rich Repeats (LRR) protein-protein interaction (PPI) domains in their N-terminal halves, while their catalytic domains are located in the C-terminal half. This catalytic portion consists of the typical combination of a Ras-of-complex (ROC) GTPase and a C-terminal of ROC (COR) domain, as well as a kinase and WD40 domains. LRRK2 has an additional Armadillo domain at the N-terminus that protrudes from the structure and is absent in LRRK1 (Myasnikov *et al*., 2021; Reimer *et al*., 2023). Both proteins exhibit similarities not only in domain organization but also in their substrate spectra as they phosphorylate members of the Rab GTPase family – albeit different ones – and thus play a role in the regulation of vesicular trafficking (Steger et al., 2016, Malik et al., 2021). Despite these similarities, there are substantial differences between the two kinases. LRRK1 is implicated in the pathogenesis of osteopetrosis (Iida et al., 2016). Since there is very limited information on its substrates or direct protein-protein interactors, it is considered a member of the “dark kinome” (Berginski *et al*., 2021). LRRK2 on the other hand has been studied extensively since its discovery in 2004 (Paisán-Ruíz *et al*., 2004; Zimprich *et al*., 2004) as it is the most frequently mutated gene in familial Parkinson’s disease (PD) (di Fonzo *et al*., 2005) and is commonly dysregulated in both idiopathic and sporadic forms of the disease (Gilks *et al*., 2005; Lesage *et al*., 2007; Nalls *et al*., 2014; Shu *et al*., 2019). Additionally, LRRK2 dysfunction has been associated with Crohn’s disease (Barrett *et al*., 2008; Hui *et al*., 2018), leprosy (Zhang *et al*., 2009; Wang *et al*., 2015), tuberculosis (Härtlova *et al*., 2018; Wang *et al*., 2018), and cancer (Saunders-Pullman *et al*., 2010; Feng, Cai and Chen, 2015; Lebovitz *et al*., 2021).

LRRK1 and LRRK2 are both serine/threonine kinases that have a DYG catalytic motif in their active site in place of the classical DFG motif (Taylor *et al*., 2020). Many of the described pathogenic variants of LRRK2 are known to have increased kinase activity (West *et al*., 2005; Kalogeropulou *et al*., 2022). Identification and investigation of LRRK2 substrates using selective inhibitors is therefore an important strategy for both gaining wider understanding of LRRK2-associated disease pathogenesis and the potential cellular impact of therapeutic inhibition. Given LRRK2’s status as a key therapeutic target, numerous high affinity and high specificity kinase inhibitors have been designed and characterized in recent years (Hu *et al*., 2023). Due to its high selectivity and potency, MLi-2 is one of the best characterized and most widely used inhibitors for the study of LRRK2 cell biology (Fell *et al*., 2015). In contrast, very few specific kinase inhibitors have been developed and validated for LRRK1. The chemical compound IN04 has previously been described as a potential LRRK1 inhibitor based on virtual homology modelling (Si *et al*., 2019). Further, its binding mode to the LRRK1 kinase active site was predicted (Chen et al., 2021). However, experimental validation of IN04’s impact on LRRK1 and its downstream substrates is lacking.

Previous mass spectrometry (MS)-based proteomics studies have investigated cellular interaction partners of LRRK2. They revealed possible roles for LRRK2 as regulator or interactor of actin cytoskeleton dynamics, vesicular trafficking, 14-3-3 proteins, and the Wnt signaling pathway among others (Meixner *et al*., 2011; Beilina *et al*., 2014; Reyniers *et al*., 2014; Steger *et al*., 2016; Salašová *et al*., 2017). For LRRK1, links to epidermal growth factor receptor (EGF-R) signaling and vesicular trafficking have been described (Hanafusa *et al*., 2011; Reyniers *et al*., 2014; Tomkins *et al*., 2018). However, to the best of our knowledge, there has been no study yet that investigated *in situ* captured cellular interaction partners for both, LRRK1 and LRRK2, and correlated them with changes in phosphorylation in presence of LRRK1 or LRRK2 targeting small molecules.

Using fast proximity labelling in combination with phosphoproteomics and data-independent acquisition (DIA)-based quantification, we characterize the cellular interactomes of LRRK1 and LRRK2. Further, we further quantify phosphorylation sites of LRRK1 and LRRK2 interacting proteins and detect abundance changes in presence of MLi-2 and IN04 – thereby identifying potentially novel interactors and substrates. Taken together, our data provide a useful resource for future studies of the cellular role and function of LRRK1 and LRRK2 and their potential use as therapeutic targets.

## Results

### Proteome-wide interaction networks for LRRK1 and LRRK2

In this study, we employed proximity labelling mass spectrometry in combination with quantitative phosphoproteomics to obtain a comprehensive dataset on cellular interactors of LRRK2 and LRRK1. We subsequently quantified the phosphorylation state of these interactors to determine how they were modulated by different LRRK-targeting compounds (**Figure 1**). To do so, we generated N-terminal TurboID-LRRK1 and LRRK2 fusion constructs containing an additional FLAG tag for biochemical enrichment (**Figure 1A**). We then used TurboID proximity labelling (Cho *et al*., 2020) in combination with a quantitative (phospho-) proteomics workflow (Zhang *et al*., 2022) based on data-independent acquisition to investigate cellular protein-protein interactions in HEK293T cells as a model system (**Figure 1B**; see **Materials and Methods** for details). After transient transfection of HEK293T cells with TurboID-LRRK1 or - LRRK2 fusion constructs or TurboID as a control (termed hereafter ‘LRRK1’, ‘LRRK2’ and ‘Control’), cells were either treated with DMSO, the LRRK1 targeting small molecule IN04 or the LRRK2 inhibitor MLi-2 (**Figure 1C**). After lysis, samples were enriched for biotinylated proteins via streptavidin beads. One part of the enriched fraction was directly processed for mass spectrometric analysis to obtain information on protein interactors of LRRK1 and LRRK2, while another part was further enriched for phosphorylated peptides to determine the phosphorylation state of these interactors. In a final fraction, phosphopeptides were directly enriched from the lysate to assess a broader impact of IN04 and MLi-2 on phosphorylation across the proteome (*see* overall workflow in **Figure 1B**).

**Figure 1.**
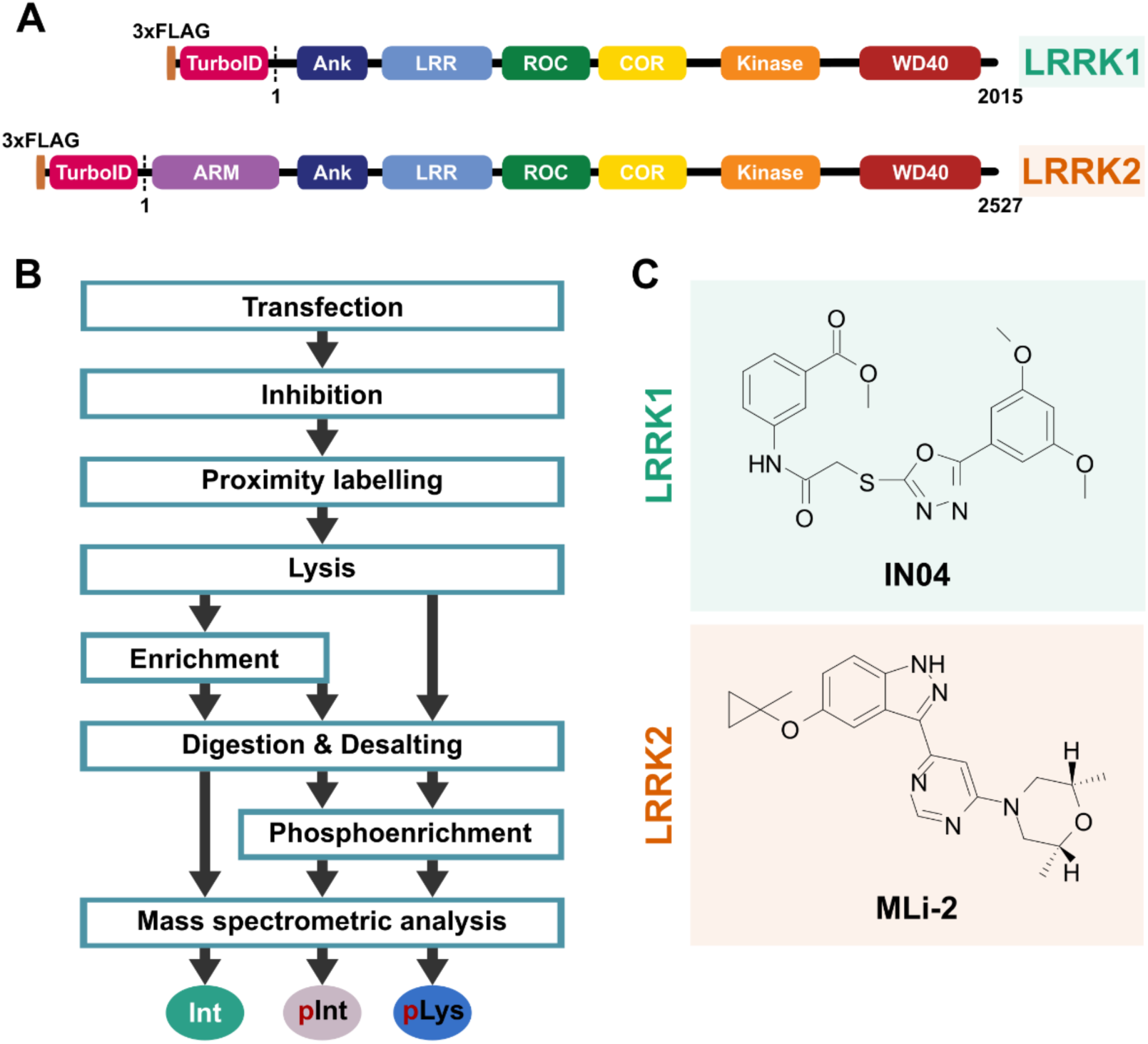
TurboID phosphoproteomics workflow to study the cellular interactomes of LRRK1 and LRRK2 and corresponding phosphorylation sites in interactors and potential substrates. (A) TurboID fusion constructs for LRRK1 and LRRK2 used in this study. Functional domains are highlighted. The start and end points of the canonical, unmodified protein sequences are indicated. (B) Overall experimental workflow for this study. HEK293T cells were transiently transfected with either TurboID-LRRK1, TurboID-LRRK2, or TurboID fusion constructs as a control. Cells were either treated with DMSO or one of the two LRRK targeting compounds (see **C**) before proximity labelling was performed. Interacting proteins were enriched via streptavidin and either directly processed for MS analysis (‘Int’, interacting proteins) or further enriched for phosphorylated peptides (‘pInt’, phosphosites of LRRK interactors). In a third fraction phosphorylated peptides were directly enriched from the lysate (‘pLys’, phosphosites of lysate proteins). (C) Chemical structures of LRRK1 (IN04) and LRRK2 (MLi-2) targeting compounds used in this study.

### Cellular interactors of LRRK1 and LRRK2 in HEK293T cells

Overall, we quantified 1,393 proteins that were significantly enriched when LRRK1 or LRRK2 were expressed compared to the control in HEK293T cells (**Figure 2A**) from a total of 6,691 identified proteins across all conditions. We validated our approach by confirming that LRRK1 and LRRK2 were present at similar levels across all replicates transfected with the respective constructs and that they were among the strongest enriched proteins (**Figure S1A and B**). Replicates expressing the respective fusion constructs were of low variance, indicating high reproducibility (**Figure S1C and D**). The potential LRRK1 and LRRK2 interactors could be grouped into five major clusters based on how they interacted with LRRK1 and LRRK2 (**Figure 2B**): proteins that were predominantly enriched for LRRK1 (clusters 1 & 2), proteins that were predominantly enriched for LRRK2 (clusters 3 & 4), and proteins that were equally enriched for LRRK1 and LRRK2 (cluster 5). For proteins that were predominantly enriched for LRRK1 or LRRK2, there were two different types of interactors. Cluster 1 and 3 contained proteins that were exclusively enriched for either LRRK1 or LRRK2, respectively, while clusters 2 and 4 consisted of proteins enriched for both LRRK proteins, but showing a stronger enrichment for one over the other.

**Figure 2.**
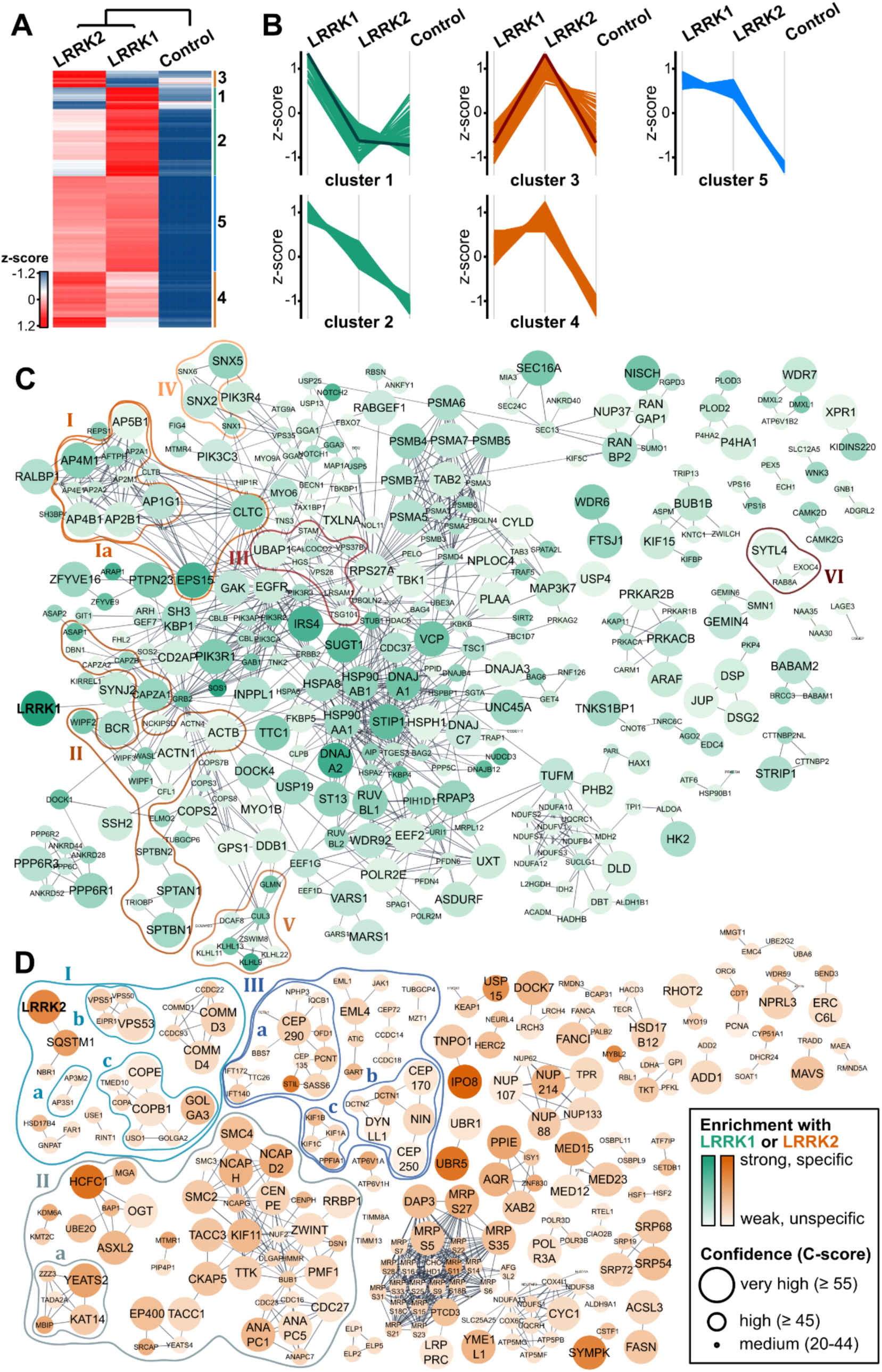
Cellular protein-protein interaction networks for LRRK1 and LRRK2. (A) Heatmap with significantly enriched interactors of LRRK1 and LRRK2 (ANOVA analysis (S0=0.1; FDR ≤ 0.01; identification in 4 of 4 biological replicates). Results are log2 transformed, z-score normalized, and biological replicates averaged. (B) Classification of enriched proteins based on their enrichment profile. Proteins preferably enriched for LRRK1 are indicated in green (cluster 1 and 2), for LRRK2 in orange (cluster 3 and 4), and proteins that were similarly enriched for LRRK1 and LRRK2 are highlighted in light blue (cluster 5). The profile graphs of LRRK1 and LRRK2 are indicated in dark green and dark orange, respectively. (C-D) Protein interaction network for the LRRK1 (in C) and LRRK2 (in D) clusters. The displayed interactions were filtered for high confidence (≥0.8; STRING) for their described interactions and further analyzed using MCL clustering. The nodes are colored continuously according to the strength of the specific enrichment according to an ANOVA and subsequent post-Hoc test (S0 = 0.1, FDR ≤ 0.01). The nodes for LRRK1 are highlighted in green and for LRRK2 in orange. The size of the nodes corresponds to the confidence of the hit identification according to the C-score of Spectronaut (see methods for details). Complexes of interest that are mentioned in the text are additionally highlighted. (C) Highlighted complexes within the LRRK1 network are (I) clathrin-coat vesicle, (Ia) AP complex proteins, (II) actin cytoskeleton, (III) ESCRT0/I complex, (IV) sortin nexins, (V) cullin3-RING ubiquitin ligase complex, (VI) Rab8A. (D) Highlighted complexes in the LRRK2 network are (I) endosomal and vesicle proteins, (Ia) AP3 complex, (Ib) EARP complex, (Ic) Golgi membrane/coated vesicle, (II) histone modifying complexes, (IIa) ATAC complex, (III) microtubular proteins, (IIIa) centrosome/cilium, (IIIb) centrosome/dynein complex, and (IIIc) kinesin complex.

To further investigate these potential interactors, we used interaction network analysis including only high confident interactions (STRING, confidence ≥ 0.8) and MCL clustering (**Figure 2C, D, and S2A**). The LRRK1 interactome of proteins preferentially associated with LRRK1 obtained from clusters 1 and 2 contained 481 proteins (**Figure 2B, C**; see **Supplementary Data**). Among the putative LRRK1 interactors were many members of the Epidermal Growth Factor Receptor (EGFR) signaling pathway (**Figure 2C**) but also a large number of proteins that are part of synaptic vesicle trafficking such as the Endosomal Sorting Complexes Required for Transport (ESCRT)-0/I complex, sorting nexins SNX1,2,5 and 6, AP2 complex proteins or clathrin-coat vesicle proteins that are all connected to early endosomal trafficking and the *trans*-Golgi network (Cui *et al*., 2022). Both the EGFR and the ESCRT-0 complex have been previously described as interactors of LRRK1 (Hanafusa *et al*., 2011; Ishikawa *et al*., 2012). Another part of this interactome network was composed of proteins that are linked to the actin cytoskeleton and its organization, corresponding to literature data, in which LRRK1-deficient cells have been described to contain deformed actin structures (Xing et al., 2017; Si et al., 2019).

The LRRK2 interactome of proteins preferentially enriched with LRRK2 expression was composed of 392 quantified proteins in total (**Figure 2A and B**; cluster 3 & 4; see **Supplementary Data**). The LRRK2 interaction network (**Figure 2D**) contained several proteins and protein complexes connected to endosomal trafficking, such as the previously described LRRK2 interaction partners SQSTM1 or the AP3 complex (AP3M2, AP3S1) (Kuwahara *et al*., 2016; Kalogeropulou *et al*., 2018) as well as Golgi and vesicle coat proteins (Biskup *et al*., 2006; MacLeod *et al*., 2013; Bae and Lee, 2020; Wei *et al*., 2023). Another interesting member of this network was the EARP (endosome-associated recycling protein) complex (Schindler et al., 2015) whose subunits Vps53, Vps51 and Vps50 were previously identified as LRRK2 interactors in a recent study which described LRRK2 as a scaffold for Soluble N-ethylmaleimide-sensitive factor attachment protein receptors (SNARE) association (Beilina et al., 2020). We also identified multiple members of the microtubule cytoskeleton as well as the centromere as potential LRRK2 interactors. Notably, this part of the network contained proteins that have been associated with cilium assembly. Finally, a number of identified potential LRRK2 interactors can be annotated to histone deacetylation, acetylation, and methylation such as the Ada2a-containing (ATAC) complex. While these interactions have not yet been associated with LRRK2, it has been previously described that LRRK2 interacts with and phosphorylates histone deacetylases such as HDAC3 (Han *et al*., 2017; Kim *et al*., 2019). Thus, our data further supports a potential role of LRRK2 in modulating epigenetic histone modification.

Among the potential interactors that were similarly enriched for both LRRK fusion constructs (**Figure 2A and B**; cluster 5) were many proteins that are connected to microtubular transport and the actin cytoskeleton as well as the Golgi apparatus and vesicle trafficking (**Figure S2A,** see **Supplementary Data**). The Rab GTPases play a vital role in vesicular trafficking and several are well established as bona fide interactors and substrates of LRRK1 or LRRK2. We confidently identified 20 Rab GTPases in our dataset and 8 of these were significantly enriched for either of the LRRK proteins (**Figure S2C and D**). Rab14, Rab8A and Rab39A were significantly enriched for both, LRRK1 and LRRK2, while Rab34 was only enriched for LRRK1 (**Figure S2C and D**). Notably, Rab8A was preferentially enriched for LRRK1 despite being a well characterized LRRK2 substrate (**Figure 2C**, complex VI)(Steger *et al*., 2016). Of note, Rab8A was significantly enriched with LRRK2 when compared directly against the control (**Figure S2D**). Moreover, other known LRRK2 substrates Rab10 and Rab12 (Steger *et al*., 2016) were also identified as LRRK2 interactors in our dataset, although not statistically significant (i.e. with a p-value ≤ 0.05). The transient nature of kinase-substrate relationships could be a possible explanation why they were not enriched to a stronger extent.

Several 14-3-3 proteins, which had previously been described as LRRK2 interactors, were also part of this cluster (Beilina *et al*., 2014; Reyniers *et al*., 2014). Finally, proteasomal subunits, chaperones and proteins of the translational machinery were also included in this cluster which could be remnants from the LRRK proteins’ life cycle.

A direct comparison of the LRRK1 and LRRK2 interactomes further demonstrated this overlap between the two kinases in terms of interacting pathways. This was particularly true for vesicular trafficking (**Figure S2A**). AP complex proteins are important throughout the vesicular transport system with different roles and localizations for the specific complexes (Sanger *et al*., 2019; Cui *et al*., 2022). While AP-3 complex proteins are in their majority specifically enriched for LRRK2, in line with literature which described the AP-3 complex as an interactor of LRRK2 at endosomes (Kuwahara *et al*., 2016), we identified AP-2 as well as AP-4- and -1 complex subunits as specifically enriched for LRRK1 (**Figure S2B**).

### Impact of LRRK targeting compounds on the phosphorylation status of LRRK1 and LRRK2 and their interactomes

Next, we asked how the interactomes and phosphorylation status of both LRRK1 and LRRK2 were impacted by the addition of LRRK-targeting compounds. To do this, we used IN04, proposed by an earlier study to inhibit LRRK1 (Si *et al*., 2019) and the well characterized, specific LRRK2 kinase inhibitor MLi-2 (Fell *et al*., 2015). Given the lack of experimental validation of IN04 as a LRRK1 kinase inhibitor, we first evaluated the effect on LRRK1 mediated phosphorylation of Rab7 at S72. Under the tested conditions, an inhibition of Rab7 pS72 was not observed (**Figure S3A**) in agreement with previous studies (Malik *et al*., 2021). Apart from inefficient or ineffective inhibition, this could be explained by the presence of other Rab7A-phosphorylating kinases such as TBK1. However, we saw a clear effect of IN04 on LRRK1 phosphosites in cells strongly suggesting that IN04 indeed affected LRRK1 phosphorylation as the presence of IN04 led to a significant downregulation of six LRRK1 phosphosites at different positions (**Figure 3B**). Here, phosphorylation at T606 exhibited the strongest effect observed with this inhibitor. This residue is located in the last of the leucine-rich repeats, while the other residues are either located in the Roc GTPase (T964, Y847 and S849), the kinase (S1236) or the WD40 domain (S1677) of LRRK1. It is important to note that contrary to LRRK2, there is little information on the autophosphorylation behavior of LRRK1 (Deng *et al*., 2011; Civiero *et al*., 2012). LRRK1 has been described to autophosphorylate, (Ishikawa *et al*., 2012) but it is so far unknown at which sites. The presence of IN04 had little effect on the interactome of LRRK1 (**Figure S4A, C**).

**Figure 3.**
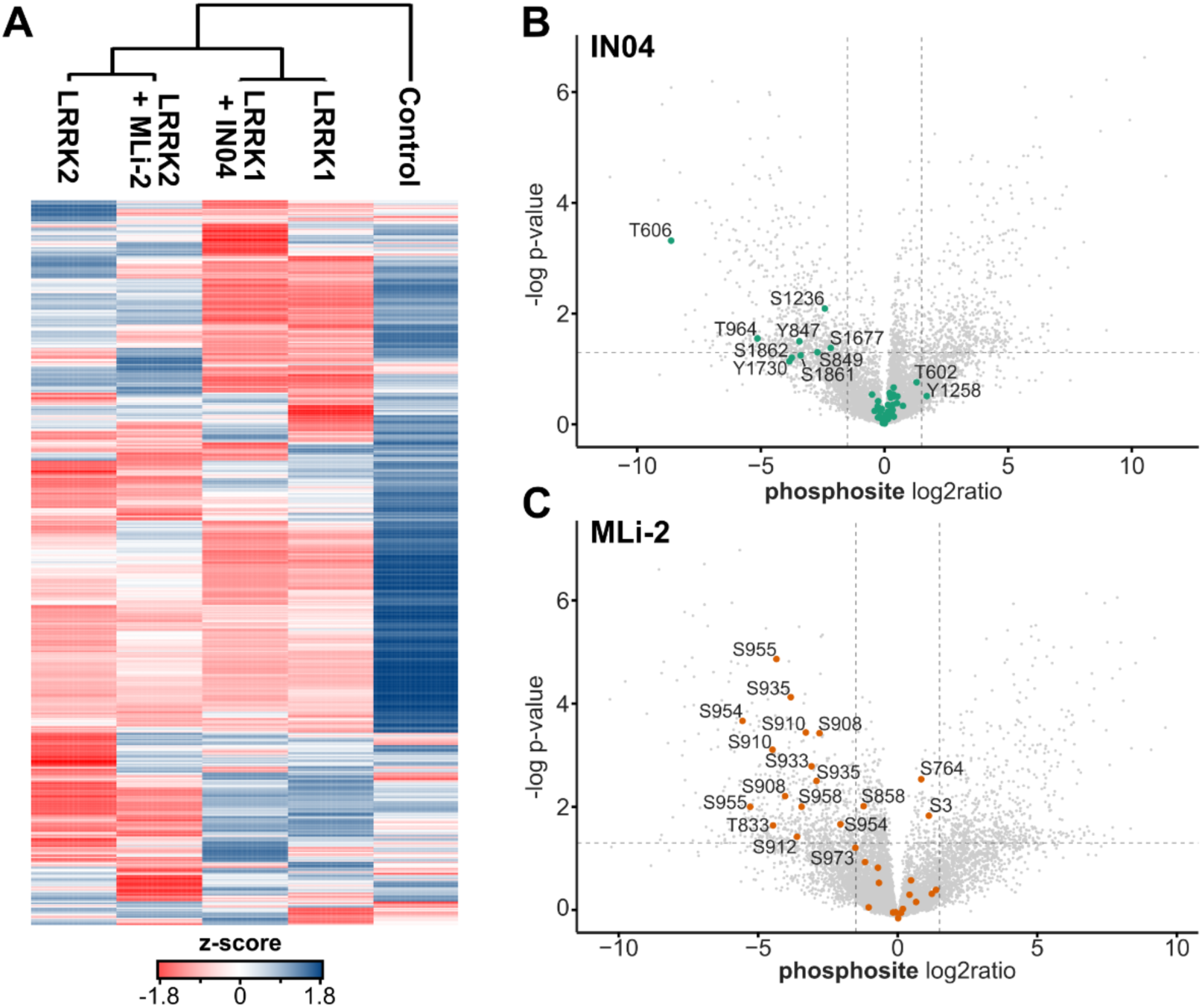
Impact of LRRK targeting compounds on the phosphorylation status of LRRK1 and 2. (A) Heatmap of significantly altered phosphosites of LRRK1 and LRRK2 interactors after addition of compounds IN04 or MLi-2 (ANOVA analysis S0=0; FDR ≤ 0.05). Results are log2 transformed, z-score normalized, and biological replicates averaged. (B-C) Volcano plots of significantly altered LRRK1 phosphosites after addition of IN04 (B) and LRRK2 after addition of MLi-2 (C). Only altered sites with a change in log2ratio ≥ |1| and p-value ≤ 0.05 (two-sample t-test) were considered significant. Phosphosites that belong to LRRK2 are indicated in orange, and to LRRK1 in green.

To evaluate a successful LRRK2 inhibition with MLi-2, we assessed phosphorylation of substrates Rab10 and Rab12 at T73 and S106, respectively, (Steger *et al*., 2016, 2017) as well as the phosphorylation of LRRK2 S935 as marker for type-I inhibition by western blotting (Tasegian *et al*., 2021). Upon addition of MLi-2, we saw a clear inhibition of phosphorylation for all three targets (**Figure S3B**). LRRK2 inhibition by MLi-2 was detectable for at least 3 hours at 1 µM. The quantification of significantly regulated phosphosites revealed that inhibition of LRRK2 by MLi-2 resulted in a significant reduction of a large number of LRRK2 phosphosites and these sites were predominantly serines residues that were located at residues 900-960 of LRRK2 (**Figure 3C**), a region which is known to harbor phosphorylation sites that are modulated by type-I inhibitors such as MLi-2 (Tasegian *et al*., 2021). Among the most strongly downregulated phosphorylation sites were S910, S935, S954 and S955 confirming literature (Dzamko *et al*., 2010; Tasegian *et al*., 2021).

Again, we saw little difference between interacting proteins of LRRK2 in the presence or absence of MLi-2 (**Figure S4B**). However, some interesting differences were distinguishable. LRRK2 interacted more strongly with proteins associated with microtubules and microtubular transport such as MID2/TRIM1 or MACF1 upon inhibition by type-I inhibitor MLi-2 (**Figure S4B**). The MID2/TRIM1 E3 ligase is a reported interactor of LRRK2 that catalyzes ubiquitination of the kinase and thereby, mediates its localization to microtubules – an effect which is reportedly increased by type-I inhibitor MLi-2 (Stormo *et al*., 2022). With respect to Rab GTPases, we saw only minor or no differences between the interactome of LRRK1 and LRRK2 in the presence or absence of their respective targeting molecule (**Figure S4C and D**). Interactions with 14-3-3 proteins on the other hand, were reduced in the presence of MLi-2 (**Figure S4C**), consistent with the fact that these proteins’ association with LRRK2 is dependent upon phosphorylation at S910 and S935 which is reduced upon kinase inhibition with MLi-2 (Dzamko *et al*., 2010; Nichols *et al*., 2010; Manschwetus *et al*., 2020).

### Impact of kinase inhibitor MLi-2 on phosphorylation state of LRRK2 interactors

In total, our phosphoproteomics analysis identified 12,148 phosphorylated sites (hereafter “phosphosites”) across all interactome conditions with high confidence, and of these 4,093 sites were significantly enriched in at least one of our four conditions, i.e., LRRK1 and 2 plus/minus addition of our two inhibitors (**Figure 3A**). We identified 65 phosphorylated sites in LRRK1 and 32 sites in LRRK2 (**Figure 3B-C**). In presence of IN04 or MLi-2, several of these phosphosites were significantly decreased as described in the previous section, while no site showed increased phosphorylation (**Figure 3B-C**).

Having validated a successful inhibition of LRRK2 phosphorylation under the experimental conditions used, we next examined how this inhibition impacted the phosphorylation status of LRRK2’s substrates and interactors.

We first assessed the phosphorylation status of the Rab GTPases as well-established substrates of LRRK2. Among the 1,386 phosphosites that were enriched directly from the lysate and significantly regulated, we were able to detect additional changes in Rab GTPase phosphorylation in absence and presence of MLi-2 (**Figure S5**). We detected a significant increase in phosphorylation of Rab12 S106 when LRRK2 (**Figure S5A**) was expressed that was significantly inhibited by MLi-2 (**Figure S5C**), while for Rab10, no phosphosites could not be identified or quantified in the lysate fraction potentially due to low protein levels. Rab12 pS106 is a previously described target for LRRK2 (Steger *et al*., 2016, 2017). This inhibition could also be confirmed by western blot (**Figure S3B**).

We then took a closer look at how MLi-2 differentially impacted the phosphoproteome of our LRRK2 interactome (**Figure 4**). We mapped phosphosites that were significantly decreased by MLi-2 onto our interactome map derived from **Figure 2D** in order to get further insight into how the inhibitor affected phosphorylation of LRRK2 interactors and potential substrates (**Figure 4**). We identified several protein complexes that were specifically associated with LRRK2, e.g., the centrosome, COMMD proteins or the mitochondrial small ribosomal subunit, that did not have phosphosites regulated by MLi-2. However, for other complexes, such as the EARP complex (Vps50), Golgi-associated proteins and centromeric proteins, we identified multiple phosphosites that were decreased in presence of MLi-2 indicating a functional relationship with LRRK2 kinase activity (**Figure 4**).

**Figure 4.**
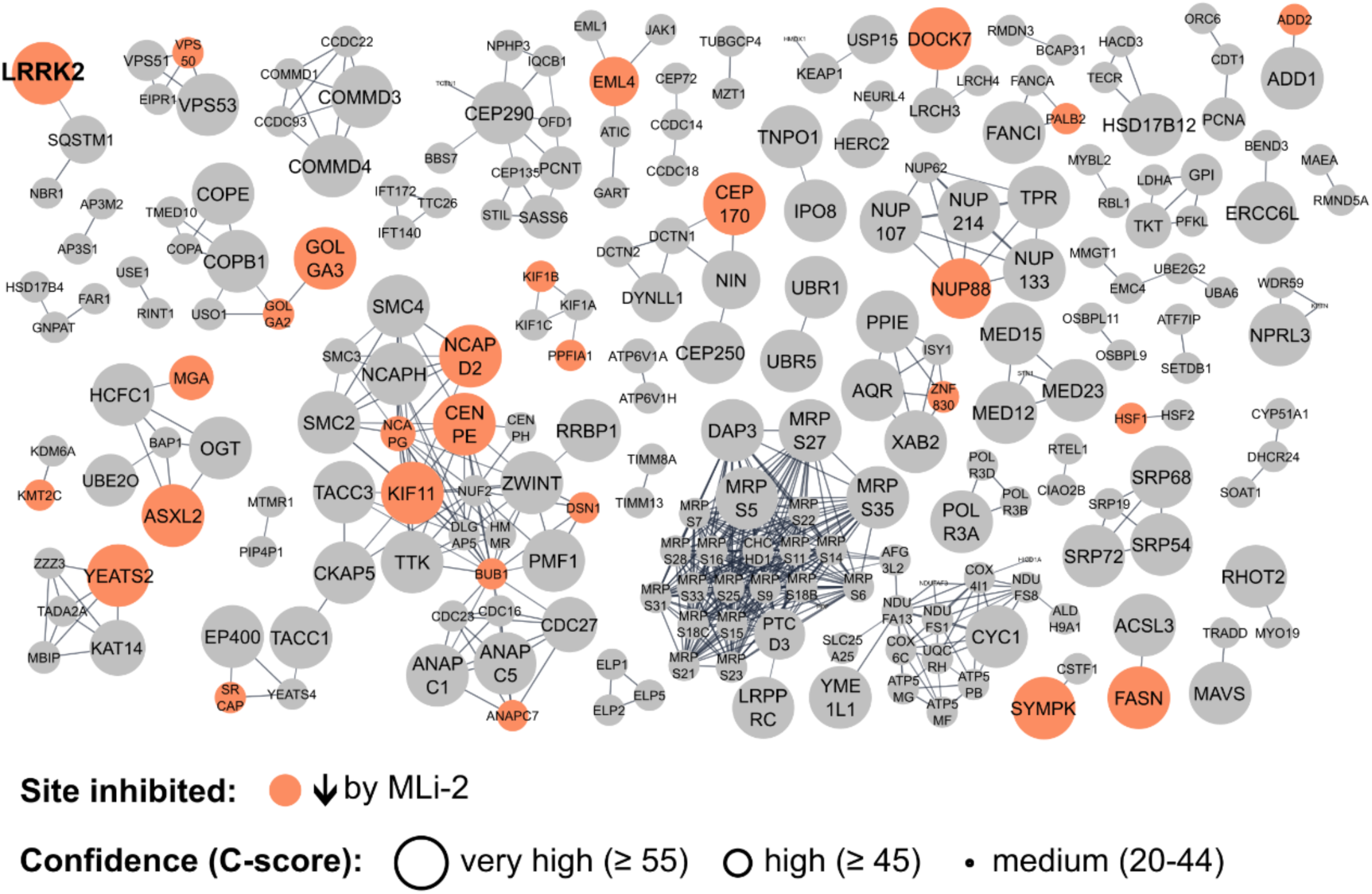
Impact of LRRK2-specific kinase inhibitor MLi-2 on LRRK2 interactors. Significantly decreased phosphosites on LRRK2 specific interactors that are inhibited by MLi-2 (i.e. clusters 3 and 4 from Figure 2B, D). Proteins with phosphosites inhibited by MLi-2 are indicated in orange. Phosphosites were regarded as significantly altered by ANOVA analysis (S0 = 0, FDR ≤ 0.05) and only sites with negatively altered phosphosites were overlayed with the interactome data.

### LRRK2 inhibition reveals two potential novel substrates

Our comprehensive approach, identifying the impact of LRRK2 kinase inhibition by MLi-2 on phosphorylation of the LRRK2 interactome, allowed us to gain information on potential LRRK2 substrates (**Figure 4**). However, to further enhance this analysis, we mapped the quantities of a specific phosphosite against the overall abundance of the respective protein, allowing us to dissect if MLi-2 inhibition impacted not only the phosphorylation state of an interactor but also affected the strength of its interaction with LRRK2 (**Figure 5**; see also workflow **Figure 1B**). We reasoned that using these combined data could be a good indicator to identify potential novel substrates. A substrate that would bind to its kinase but not be phosphorylated due to inhibition of kinase activity might not be released as fast, as suggested in the literature (Adams, 2001; Sommese and Sivaramakrishnan, 2016) and thus, appear as a stronger enriched interactor in our data. At the same time, the phosphorylation itself would be inhibited and the respective phosphosite found to be decreased in quantity. Mapping protein and phosphosite quantities against each other also allowed us to distinguish between phosphosites that were less abundant as an effect of the inhibition or rather because the respective protein itself interacted less with LRRK2.

**Figure 5.**
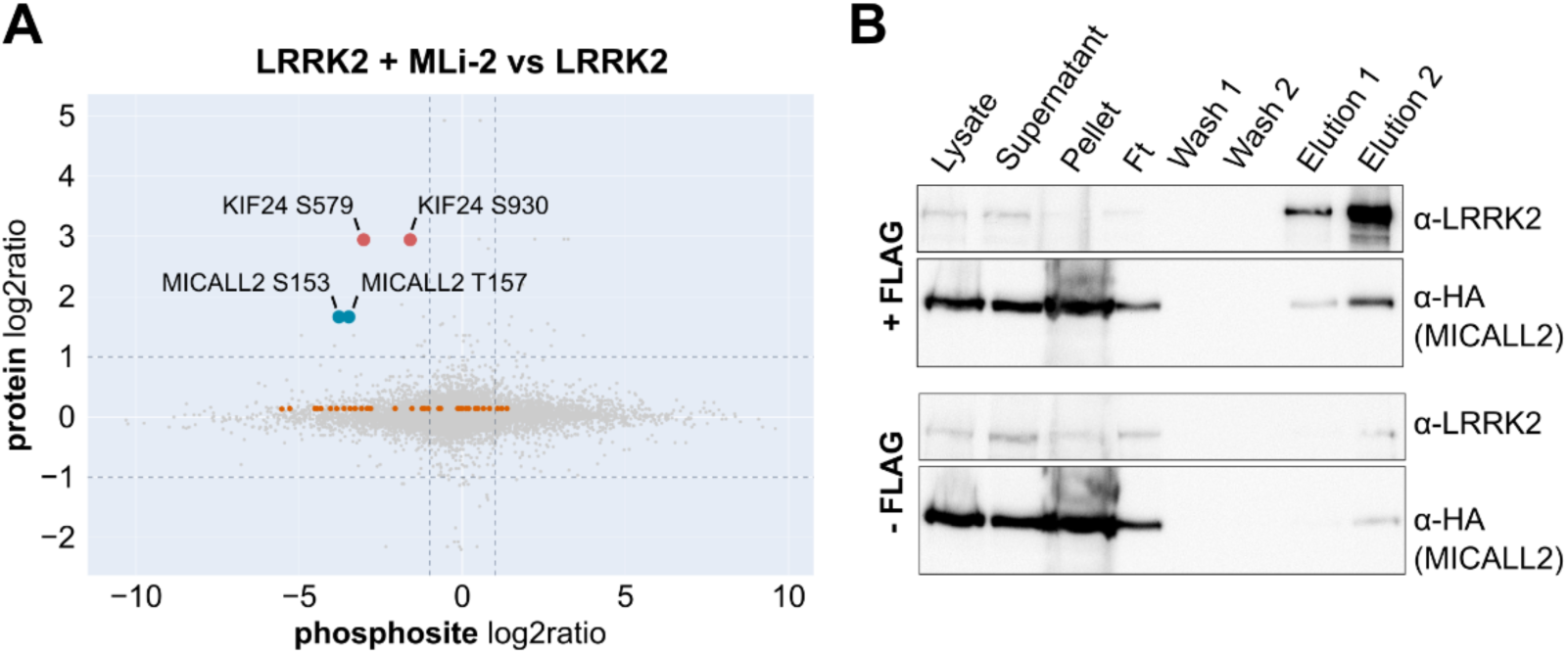
Kif24 and MICALL2 are potential novel substrates of LRRK2. (A) Graph showing the abundance change of interactome-wide significant phosphosites (x-axis) against the abundance change of the respective protein (y-axis) upon inhibition of LRRK2 by MLi-2. Novel interactors and potential substrates MICALL2 and Kif24 are highlighted in blue and red, respectively. Phosphosites identified within LRRK2 are indicated in orange as a reference. Only phosphosites are shown (p-value ≤ 0.05). (B) Co-immunoprecipitation of 3xFLAG-turboID-LRRK2 (upper panel) or turboID-LRRK2 (lower panel) and C-terminally HA-tagged MICALL2. Both constructs were transiently co-transfected into HEK293T cells. After incubation of the lysate with the anti-FLAG resin, beads were washed with lysis buffer (Wash 1) and PBS (Wash 2). Then, bound proteins were eluted with 3xFLAG peptide (Elution 1) before residual protein was eluted under harsh conditions (Elution 2, see Methods). Transfection efficiency was controlled by β-galactosidase co-transfection and sample inputs normalized thereby. Western blot representative of 2 independent experiments.

We identified two proteins, MICALL2 and Kif24, that stood out in this analysis, because their phosphosites were not only inhibited in the presence of MLi-2 but at the same time their corresponding protein level was stronger with LRRK2 (**Figure 5A**). MICALL2 is a protein involved in vesicular transport and cytoskeletal organization that has been reported to interact with Rab8 and Rab13 (Terai *et al*., 2006; Yamamura *et al*., 2008). It is also reported as one of the strongest interactors of Rab8 T72A, a non-phosphorylatable variant of Rab8 (Steger *et al*., 2016). Kif24 is a kinesin motor protein that associates with microtubules and negatively regulates ciliogenesis (Kobayashi *et al*., 2011) has recently been implicated in late-onset PD (Gialluisi *et al*., 2021).

To investigate if MICALL2 is indeed a potential novel interactor of LRRK2, we performed co-precipitation experiments using the FLAG-tag of our LRRK2 construct. We transfected HEK293T cells with our LRRK2 plasmid as well as a plasmid containing C-terminally HA-tagged MICALL2 (**Figure 5B**). We found that MICALL2 was specifically bound to LRRK2 and co-eluted with it from the anti-FLAG resin (**Figure 5B**). Thus, we concluded that there is strong evidence that MICALL2 could be a novel interactor of LRRK2. Future studies will be needed to elucidate LRRK2’s interaction with MICALL2 as well as with cilia-associated Kif24 in more detail to gain insights into how these interactions could play a role in LRRK2-associated disease mechanisms.

## Discussion

Almost ten years have passed since members of the small Rab GTPase family have been identified as LRRK2 substrates using a comprehensive phosphoproteomics-based MS approach (Steger *et al*., 2016, 2017). In addition to these Rab GTPases, only few other proteins have been identified as bona fide LRRK2 substrates, including HDAC3 or p53 (Ho *et al*., 2015; Han *et al*., 2017). However, LRRK2 is known to interact with multiple cellular compartments (Usmani, Shavarebi and Hiniker, 2021), posing the question of how many substrates are yet to be discovered. For LRRK1, far fewer interactors and substrates have been identified so far (Hanafusa et al., 2011, 2015; Kedashiro et al., 2015; Zeng et al., 2016;Reyniers et al., 2014; Tomkins et al., 2018).

In this study, we employed proximity labelling mass spectrometry using TurboID in combination with quantitative phosphoproteomics to obtain a comprehensive dataset of cellular interactors of LRRK1 and LRRK2 and their phosphorylation state in a model cell line. We then used this dataset to characterize the impact of small molecules targeting LRRK1 and LRRK2 on the phosphorylation state of the interactome and wider proteome, and to identify potentially novel substrates for LRRK2.

By fusing LRRK1 or LRRK2 with the promiscuous TurboID biotin ligase (Cho *et al*., 2020), we identified hundreds of interaction partners for these two ROCO proteins. Identification of known interaction partners such as EGFR and Grb2 for LRRK1 and SQSTM1, AP3 complex proteins, Rab8A for LRRK2 validated our approach and demonstrated its ability to capture previously identified interactors (**Figure 2, S2**). Of note, Rab8A was found as an interactor of LRRK2 (**Figure S2D**) as well as LRRK1 (**Figure S2C**) and was enriched more strongly with LRRK1 (cluster 2 from **Figure 2B** and **C**). LRRK1 is known to be essential for endosomal trafficking of the EGFR from the plasma membrane to late endosomes (Hanafusa *et al*., 2011; Ishikawa *et al*., 2012). Our extensive interaction map for LRRK1 (**Figure 2C**) includes interesting novel candidate interactors such as clathrin, or AP-2 and -4 complex proteins that could shed light on LRRK1’s role in this pathway. Further, a cluster of Sorting nexins (SNX1,2,5 and 6) which are components of the SNX-BAR complex (Seaman, 2012; Hanley and Cooper, 2021) was also identified as interactors of LRRK1. As one of the retromer subcomplexes, these proteins are responsible for protein transport between late endosomes and the Golgi and sorting of the EGFR to lysosomes (Yang *et al*., 2022) further substantiating LRRK1’s role in EGFR endosomal trafficking. Another interesting interactor in our dataset is Vps35, a component of the second retromer subcomplex (Haft *et al*., 2000). The known LRRK1 substrate Rab7a has previously been described to interact with Vps35 and to regulate the association of the Vps retromer complex with the endosome as well as to interact with PI3K, which was also present in our LRRK1 interactome map (**Figure 2C**)(Stein *et al*., 2003; Rojas *et al*., 2008).

LRRK2 has been described as interactor of many pathways in the cell among them vesicle trafficking and sorting at the Golgi (Erb and Moore, 2020).Our LRRK2 interactome (**Figure 2D**) contained Vps proteins Vps50, Vps51 and Vps53 that are components of the EARP complex, which is responsible for endocytic recycling (Schindler *et al*., 2015). Previous data had indicated an interaction of LRRK2 with this complex, although follow-up experiments suggested that LRRK2 might instead rather interact with the closely related GARP complex (Beilina *et al*., 2020). The identification of Vps50 as potential LRRK2 interactor suggests an additional or alternative interaction of LRRK2 with the EARP complex. Our data contained other proteins responsible for endocytic trafficking such as the AP3 complex, or COPI-coat vesicle transport proteins at the Golgi apparatus (COPE, COPB1, GOLGA3, GOLGA2) detailing LRRK2’s interaction with the Golgi. Another set of interesting interactors was associated with the kinetochore and histone modifications, such as the ATAC complex (Nagy *et al*., 2010). Some of the identified interactors, as the ATAC complex, have also been described to regulate tubulin acetylation (Nagy *et al*., 2010; Orpinell *et al*., 2010). This is interesting as LRRK2 has been described to preferentially associate with deacetylated microtubules, and it is known that an increase in microtubule acetylation can reverse axonal transport problems caused by pathogenic LRRK2 variants (Godena *et al*., 2014). Another study suggested that LRRK2 actively regulates the deacetylation of microtubules (Law *et al*., 2014). The deacetylation of histones was also proposed as a mechanism of how LRRK2 pathogenic variants confer neurotoxicity (Han *et al*., 2017). Taken together, our data validates known interactors of LRRK2 and identifies a series of potential novel interactors which enhances our understanding of its role in vesicle trafficking at the Golgi. It also supports a role of LRRK2 in the regulation of histone modifications that was described previously (Han *et al*., 2017). Follow up studies should focus on validating these interactions and their functional significance.

Our study also showed a significant overlap in proteins that were similarly enriched with both LRRK proteins. Previous studies had already indicated a certain overlap between the interactors of the two homologues (Reyniers *et al*., 2014; Tomkins *et al*., 2018). As the two ROCO kinases have a similar domain structure, they have even been suggested to have redundant functions and be able to compensate a loss of function for each other (Biskup *et al*., 2007; Westerlund *et al*., 2008; Tong *et al*., 2012). It is for example known that both kinases are involved in vesicle transport (Malik *et al*., 2021). Some of these previously identified interactors were also identified in our study, such as NEK1 or TUBA1B (Beilina *et al*., 2014; Reyniers *et al*., 2014; Tomkins *et al*., 2018) but we also identified proteins and complexes that have not been described for both LRRK proteins so far (**Figure S2A**). In our study, the interactors that were identified for both LRRK proteins contained many microtubule-associated proteins such as the dynein complex or the FHF complex. Interestingly, among these common interactors we have identified also regulators of ciliogenesis, such as the previously mentioned NEK1 as well as CEP97 and CCP110. All eight subunits of the conserved oligomeric Golgi (COG) complex were also part of this common interactome. The COG complex is an integral part of the Golgi apparatus and functions in the retrograde transport of endosomes to the Golgi as well as retrograde intra-Golgi trafficking (Ungar *et al*., 2002, 2006). Among others, it interacts with Rab proteins such as Rab39 or Rab10 (Willett, Ungar and Lupashin, 2013). While most members of the EARP/GARP complex were identified in the LRRK2 interactome, Vps52 was found to be similarly enriched with LRRK1 in our study. Vps52 was recently described as a direct interactor of LRRK2 (Beilina *et al*., 2020). Therefore, our data might shed light on a shared role for LRRK1 and 2 in the retrograde transport of endosomes to the Golgi.

Interestingly, many members of the 14-3-3 protein family were also identified as interactors of both LRRK1 and 2 in our study. 14-3-3 proteins have previously been described as interactors for LRRK2 but have not yet been connected to LRRK1 (Beilina *et al*., 2014; Reyniers *et al*., 2014; Tomkins *et al*., 2018). However, members of the 14-3-3 protein family have been described to bind to the known LRRK1 interactor SOS1 in its phosphorylated state (Saha *et al*., 2012). 14-3-3 proteins are interaction hubs with hundreds of known binding partners and regulate key cellular pathways (Obsilova and Obsil, 2022) and interact with specific phosphopeptide motifs (Yaffe *et al*., 1997). Our data shows that members of the 14-3-3 protein family were not only able to interact with phosphorylated LRRK2 (Manschwetus *et al*., 2020) but also with phosphorylated LRRK1. Moreover, our data supports that this interaction of 14-3-3 protein family members with LRRK2 specifically depends on the phosphorylation of serines 910 and 935 within LRRK2, as has been previously suggested in the literature (Dzamko *et al*., 2010; Nichols *et al*., 2010; Manschwetus *et al*., 2020). Consistently, our data demonstrates that the binding of 14-3-3 family members decreases when LRRK2 is inhibited by MLi-2, which results in a loss of phosphorylation at 14-3-3 binding sites (**Figure S4**). On the other hand, we saw that MLi-2 inhibition increases LRRK2’s interaction with microtubular proteins (**Figure S4B**) which is consistent with previous observations that LRRK2, once it is inhibited by type-I inhibitors, binds stronger to microtubules (Dzamko *et al*., 2010; Blanca-Ramírez *et al*., 2017; Schmidt *et al*., 2019; Deniston *et al*., 2020).

The integration of LRRK2 interacting proteins and their respective phosphosites revealed which proteins and complexes were targeted by LRRK2 inhibition and could therefore potentially consist of LRRK2 substrates (**Figure 4, 5, S5**). Our data showed that some complexes like the centrosome-related proteins contained no regulated sites indicating that they are likely interactors but not substrates of LRRK2. Other interacting complexes, such as the EARP complex, the condensing complex or kinetochore proteins DSN1 and CENPE had phosphorylated sites that were targeted by MLi-2, suggesting that they might indeed also contain potentially novel substrates of LRRK2.

A more detailed analysis of LRRK2 inhibition combining identified phosphosites with protein abundance revealed two interesting substrate candidates: Kif24 and MICALL2 (**Figure 5**). Both proteins were slightly stronger enriched after inhibition by MLi-2 while their phosphosites were significantly decreased. MICALL2 is an interactor of small Rab GTPases like Rab8 and Rab13 (Terai *et al*., 2006; Yamamura *et al*., 2008) and has been described to be important in the tubulation of recycling endosomes in concert with Rab8A, another interactor of LRRK2 (Steger *et al*., 2016; Sakane *et al*., 2021). We validated this interaction using an orthogonal co-immunoprecipitation approach (**Figure 5B**), thereby corroborating our integrated MS-based approach. Future work should investigate the functional relationship between LRRK2 and MICALL2 as well as our second hit Kif24 in more detail. For these follow-up experiments it would be advantageous to encompass a more disease-relevant cell type, or even patient-derived induced pluripotent stem cells that can be differentiated into microglia, astrocytes, macrophages, or other cell types relevant to LRRK2-associated PD. This could give rise to additional, more physiologically relevant information that might be missing in our model cell line data.

Taken together, in this study we have employed proximity labelling mass spectrometry in combination with quantitative phosphoproteomics to obtain the cellular interactome and its phosphorylation state of LRRK1 and LRRK2 in a model cell line. We then used this data to characterize the impact of kinase inhibition on the wider proteome and identified potentially novel substrates for LRRK2. Taken together, our data provide a powerful resource for future studies of the cellular role and function of LRRK1 and 2 and their potential use as therapeutic targets.

## Acknowledgements

This research was funded by Aligning Science Across Parkinson’s (grant ASAP-000519 to S.R.P., S.K. and F.S) through the Michael J. Fox Foundation for Parkinson’s Research (MJFF). This work was also supported by funding of the German Research Foundation (DFG, F.S.: 496470458 and 516836828). We thank Doreen Herzog and Andreas Marx for providing the HEK293T cell line. For the purpose of open access, the author has applied a CC BY public copyright license to all Author Accepted Manuscripts arising from this submission.

## Conflict of interest

There is no conflict of interest

## DATA AND CODE AVAILABILITY

The data, protocols and key lab materials used and generated in this study are listed in a Key Resources Table alongside their persistent identifiers in the Materials and Method section. The mass spectrometry proteomics data have been deposited to the ProteomeXchange Consortium via the PRIDE partner repository (Perez-Riverol *et al*., 2022) with the dataset identifier PXD067057 (Token: oBMf19A5vZpk; Username; Password: 4ZCpKPoygWDs).

## Material and Methods

### Key Resources Table

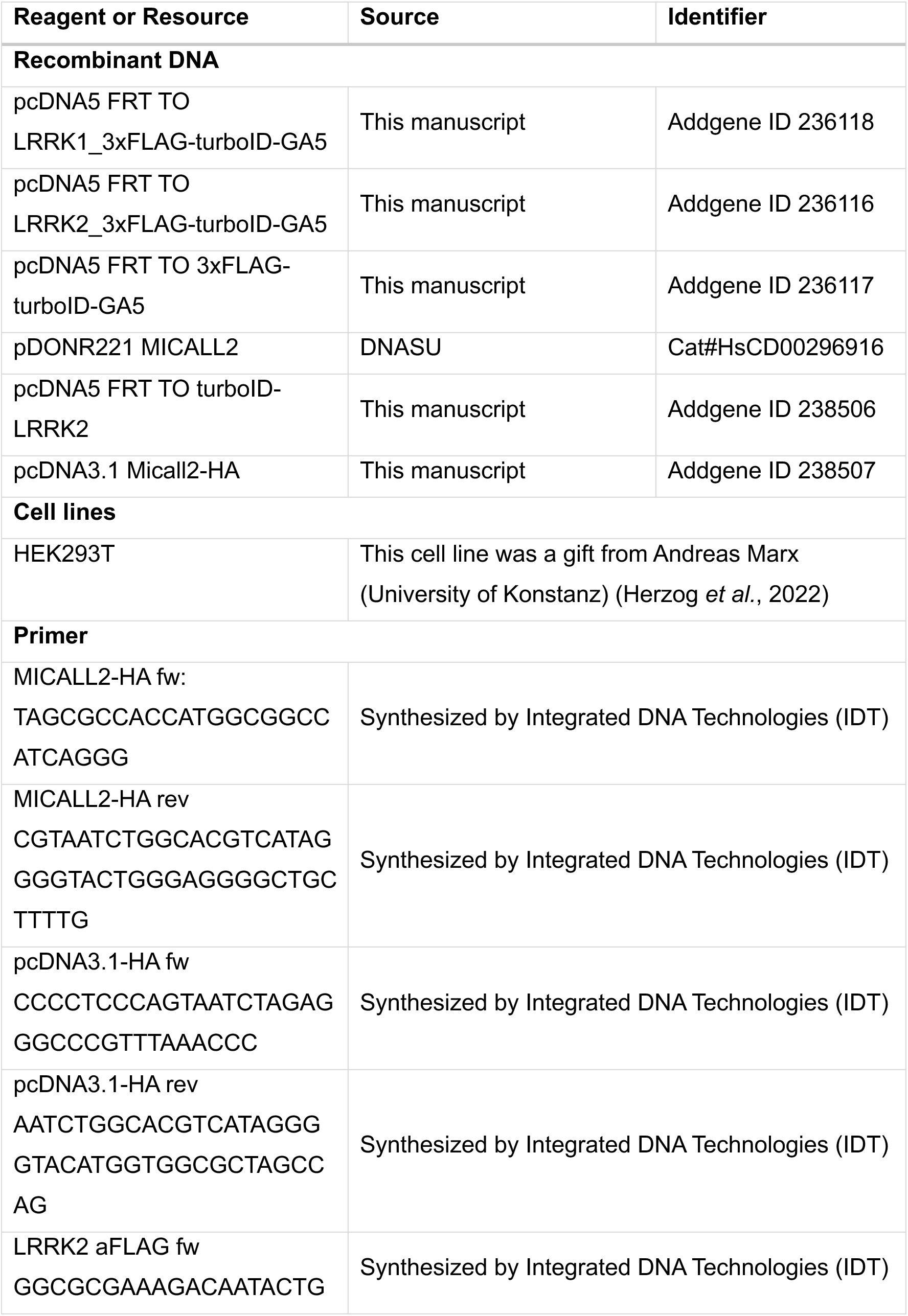
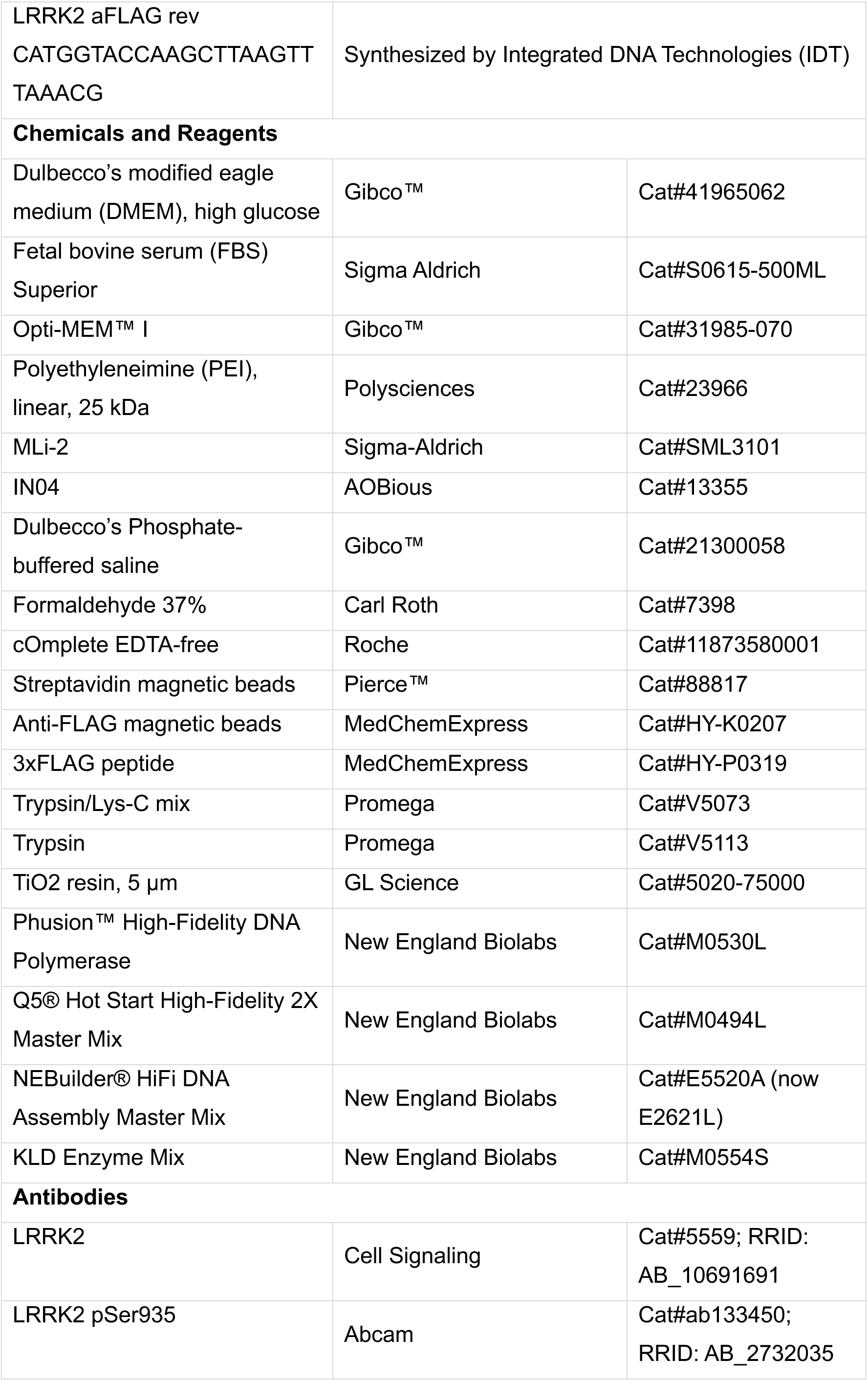
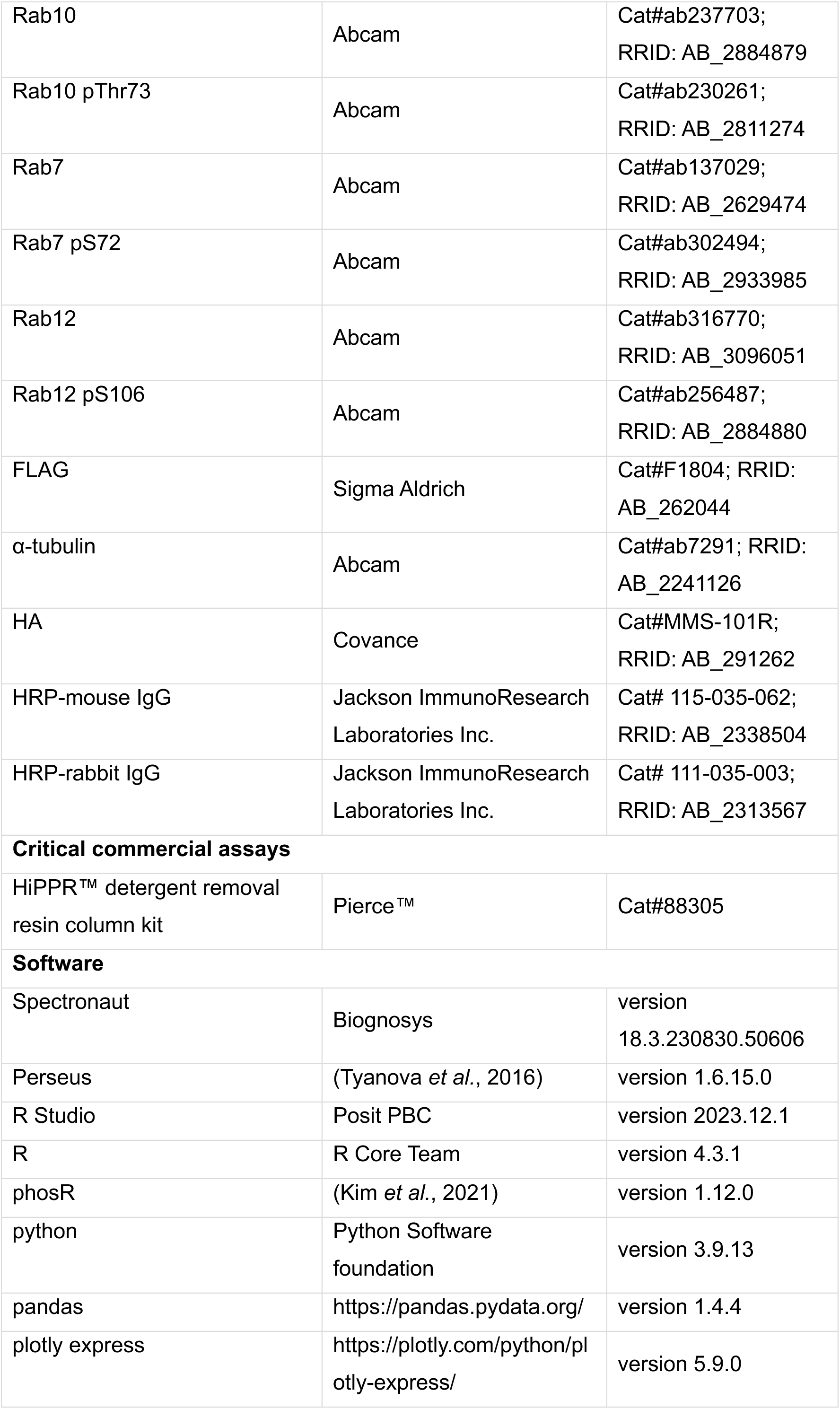
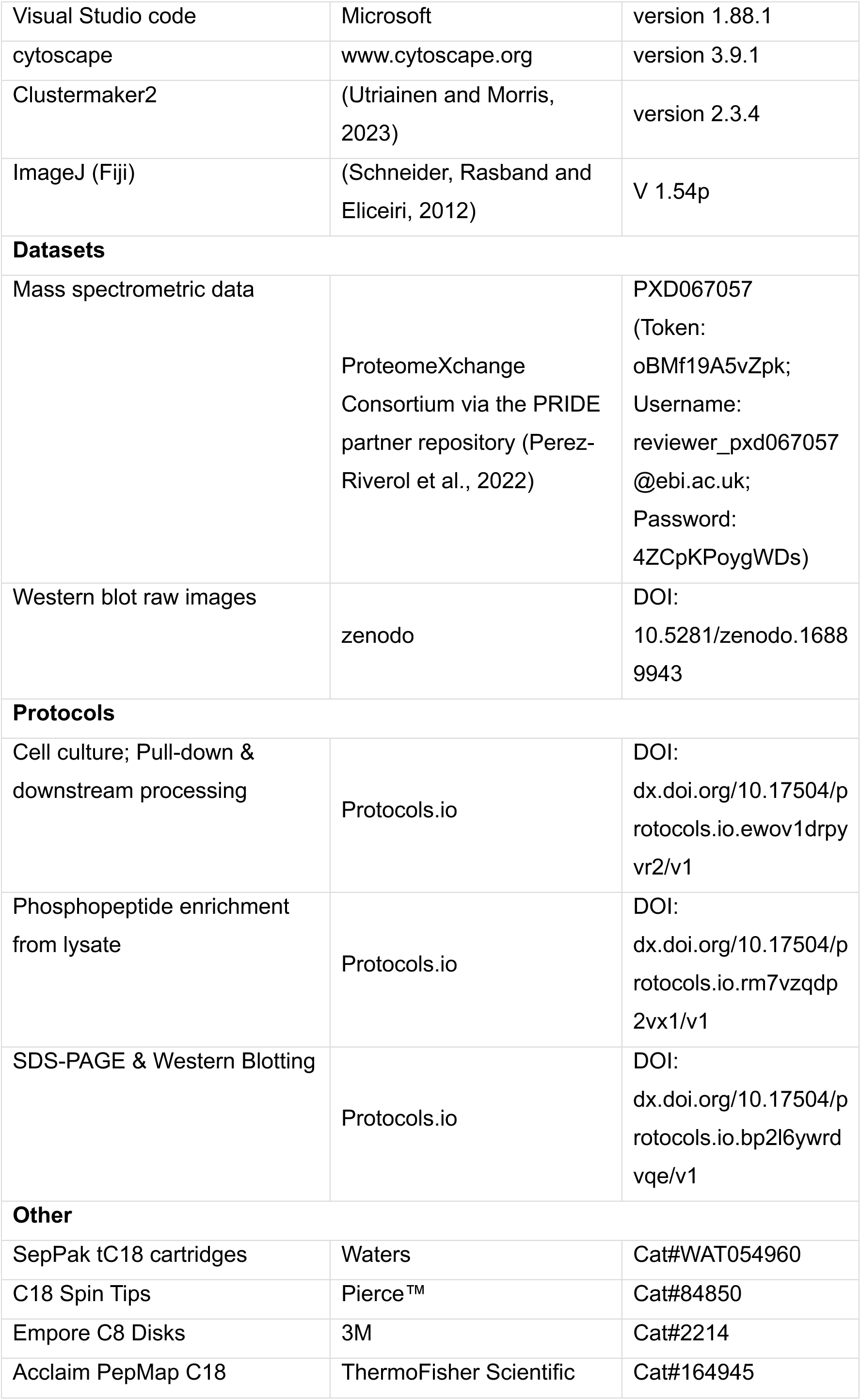

### Cell culture

HEK293T cells were grown in Dulbecco’s Modified Eagle Medium (DMEM) with 10 % fetal bovine serum (FBS) at 37 °C, 95 % humidity and 5 % CO_2_. For transient transfection, cells were grown in 15 cm dishes to 50-60 % confluency before fresh medium was added prior to transfection. For one replicate, four 15 cm dishes were transfected with the same transfection mix. Plasmid was mixed with polyethyleneimine (PEI) in OptiMEM, incubated for 15 min at room temperature before it was added dropwise to the cells. After 24 h, the medium was exchanged, and cells were ready for TurboID experiments.

### Molecular Cloning

Human MICALL2 was amplified from pDORN211 vector with additional YPYDV coding sequence part for C-terminal HA tag via PCR using Phusion™ High-Fidelity DNA Polymerase. pcDNA3.1 backbone was amplified with the additional PDYA coding sequence peart for the HA tag PCR utilizing Q5® Hot Start High-Fidelity 2X Master Mix. PCR products were assembled via NEBuilder® HiFi DNA Assembly Master Mix to pcDNA3.1 Micall2-HA vector.

3xFLAG tag was removed via Phusion™ PCR to amplify pcDNA5 FRT TO LRRK2_turboID-GA5 vector. The PCR product was ligated utilizing KLD Enzyme Mix. All generated constructs were sequenced from GENEWIZ from Azenta Life Sciences.

### MICALL2 co-immunoprecipitation

For transient co-transfection, cells were grown in 15 cm dishes to 50-60 % confluency before fresh medium was added prior to transfection. For each replicate, two 15 cm dishes were transfected with the same transfection mix, containing respectively 7.5 µg pcDNA5 vector for LRRK2 transfection with and without 3xFLAG, 7.5 µg pcDNA3.1 vector for MICALL2 transfection and 1 µg pcDNA3 vector for β-galactosidase transfection. Plasmids were mixed with PEI in OptiMEM, incubated for 15 min at room temperature before the mixture was added dropwise to the cells. 24 h after transfection, Cells were washed twice with warm 1x PBS, incubated with 0.025 % formaldehyde in PBS for 10 min, quenched with 50 mM Tris pH 7.4 and then harvested in ice-cold 1x PBS. Cell pellets were washed twice in PBS before they were snap-frozen and stored at -80 °C.

After thawing on ice, cell pellets were resuspended in 8 mL NP-40 buffer (25 mM Tris-HCl pH 7.4, 150 mM NaCl, 1 % NP-40, 5 mM MgCl2, 1x cOmplete EDTA-free, 1 µg/mL Aprotinin/Leupeptin, 100 µM Pefablock), incubated for 15 min at 4 °C and centrifuged at 15,000 g for 20 min at 4 °C. Transfection efficiency was controlled via β-galactosidase assay and equilibrated anti-FLAG magnetic beads (20 µL slurry per replicate) were added prior incubation for 20 h at 4 °C with gentle agitation. The beads were pelleted at 2000 g for 15 min at 4 °C and the supernatant carefully decanted using a magnet. 1 mL NP-40 buffer was used to transfer the beads to a 1.5 mL vial and washed twice with 1 mL NP-40 buffer and twice with 1x PBS by using a magnetic rack. Finally, the proteins were eluted by adding 2x 10 µL elution buffer 1 (1 mg/mL 3× Flag-Peptide, 50 mM Tris, 0.15 M NaCl, pH 7.4) with gentle shaking for 10 min at rt. Elution 1 was mixed with Laemmli sample buffer, and remaining beads were heated in Laemmli buffer as elution 2. Then, both elutions were analyzed by SDS-PAGE and Western Blotting.

### SDS PAGE and Western Blotting

For SDS-PAGE and Western Blot experiments, cells were seeded and transfected in 6-well dishes unless described otherwise. Cells were lysed in RIPA buffer (50 mM Tris pH 8, 150 mM NaCl, 0.5 % sodium deoxycholate, 0.1 % SDS, 1 % NP-40, 1 mM DTT, 1x EDTA-free cOmplete) containing phosphatase inhibitors (5 mM NaF, 5 mM β-glycero phosphate, 1 mM Na_3_VO_4_) for 20 min at 4 °C and cell debris was pelleted by centrifugation at 16,000 g for 10 min and 4 °C. Protein concentration of the supernatant was determined by bicinchoninic acid assay and results used to normalize input for analysis by SDS-PAGE. Samples were mixed with 2x Laemmli sample buffer (62.5 mM Tris-HCl pH6.8, 100 mM DTT, bromophenole blue), loaded onto a 7.5, 10 or 12.5 % SDS-PA gel and resolved for 80 min at 80-110 V. Proteins were transferred onto a 0.22 µm or 0.45 µm (for proteins > 50 kDa) PVDF membrane by Western Blotting in transfer buffer (12.5 mM Tris-HCl pH 8.3, 100 mM glycine) for 90 min at 60 V. The membrane was blocked by 5 % milk in TNE-T buffer, incubated with the primary antibody in TNE-T or 5% BSA in TNE-T for phosphosite antibodies for 16-20 h at 4 °C before it was incubated with the secondary antibody for 1.5 h at 21 °C. The membrane was treated with detection reagent to detect chemiluminescence by LAS-3000. For detection of Rab protein levels after phosphosite detection, membranes were stripped of bound antibodies with stripping buffer (20 mM Tris-HCl pH 7.4, 6M guanidine hydrochloride, 0.02% NP-40, 0.1M β-mercaptoethanol) for 10 min, re-blocked, and treated as described before. Images were processed by adjusting brightness and contrast for better visibility of signals using ImageJ (Schneider, Rasband and Eliceiri, 2012).

### TurboID proximity labelling

After transient transfection, 1 µM MLi-2, 5 µM IN04 in DMSO or DMSO as control were added to the transfected cells and incubated for 1 h at 37°C, before 50 µM biotin in DMSO (100 mM stock) was added and incubated for another hour at 37 °C. Cells were washed twice with warm 1x PBS, incubated with 0.025 % formaldehyde in PBS for 10 min, quenched with 50 mM Tris pH 7.4 and then harvested in ice-cold 1x PBS. Cell pellets were washed twice in PBS before they were snap-frozen and stored at -80 °C.

### Lysis and streptavidin enrichment

After thawing on ice, cell pellets were resuspended in 16 mL lysis buffer (25 mM Tris-HCl pH 7.4, 150 mM NaCl, 1 % NP-40, 5 mM MgCl2, 1 mM Na_3_VO_4_, 5 mM NaF, 5 mM β-glycero phosphate, 1x cOmplete EDTA-free, 1 mM DTT), incubated for 15 min at 4 °C and centrifuged at 16,000 g for 20 min at 4 °C. After equilibrating Streptavidin magnetic beads in buffer, they were added to the supernatant (5 mg beads per sample) and incubated for 20 h at 4 °C with gentle agitation. The beads were pelleted by centrifugation at 2,000 g for 15 min at 4 °C and the supernatant carefully decanted using a magnet. 1 mL NP-40 buffer was used to transfer the beads to a 2 mL vial and washed twice with 1 mL NP-40 buffer by using a magnetic rack. To prepare them for on-bead digestion, beads were equilibrated with 2x 1 mL 50 mM ammonium bicarbonate (AB).

For the third fraction in which phosphopeptides were directly enriched from cell lysate, 200 µL of lysate were removed and proteins precipitated by adding 6 volumes of ice-cold acetone. Then, samples were incubated for 15 min at -80 °C and for 90 min at -20 °C before they were centrifuged for 10 min at 16,000 g and 4°C. The acetone containing supernatant was removed and pellets dried at room temperature.

### Digestion and peptide processing

After equilibration with ammonium bicarbonate, beads were resuspended in 150 µL 8 M urea, before they were incubated with 3 mM TCEP for 30 min at 37 °C and subsequently, with 6 mM iodoacetamide (IAA) for 30 min at 23 °C in the dark to reduce and alkylate cysteines. For digestion, urea concentration was diluted to 4M using 50 mM AB, then, 3 µg Trypsin/Lys-C mix were added, and Lys-C digestion was allowed for 3.5 h at 37 °C. Afterwards, urea was further diluted to 1 M, additional 2 µg trypsin added, and trypsin digestion performed for 18 h at 37 °C. To remove residual detergents that would interfere with mass spectrometric analysis, the samples were incubated with HiPPR detergent removal resin (200 µg resin per sample) for 10 min, eluted and then acidified by adding formic acid to a final concentration of 2 % to stop digestion completely. Then, acetonitrile (ACN) was added to a final concentration of 1 % and peptides were desalted using Sep-PAK C18 cartridges. Cartridges were wetted with pure ACN, equilibrated twice with 1 % ACN, 0.1 % formic acid, before samples were slowly applied to the columns. Then, the bound peptides were washed twice with 1 % ACN, 0.1 % formic acid and eluted with 50 % ACN, 0.1 % formic acid. For further analysis, samples were split so that 90 % could be used as input for phosphoproteomics enrichment of the interaction fraction and 10 % for peptide analysis of interacting proteins. Precipitated samples from the third fraction described in the previous section, were treated similarly as the on-bead samples, i.e. protein pellet was dissolved in 150 µL 8M urea for 15 min at 37 °C and 850 rpm before reduction, alkylation, digestion and further peptide processing was done. All three fractions were dried and stored at -20 °C until further analysis.

### Phosphopeptide enrichment

Approximately 50 µg of dried peptides for the interactome and 200 µg for the lysate fraction were resuspended in 200 µL Loading Buffer 1 (0.1M glycolic acid, 70 % ACN, 5 % TFA), while TiO2 beads were prepared by resuspension in Loading buffer 1 (1 mg beads in 10 µL buffer) and subsequent incubation at 1200 rpm for 20 min at 22 °C. Equilibrated beads were added to the peptides at a peptide:bead ratio at 1:10 and incubated for 20 min at 22 °C and gentle agitation. In the meantime, self-made, single-layered C8 Stage tips were prepared (Rappsilber, Mann and Ishihama, 2007) and equilibrated with Loading buffer 1. Incubated beads were settled by centrifugation at 10 000 g for 2 min and the supernatant carefully transferred to a fresh low binding tube for a second enrichment step. Residual supernatant and the beads were added to the equilibrated C8 Stage tips, and centrifuged at 500 g for 2 min. Then, beads were washed with 50 µL Wash buffer 1 (80 % ACN, 1 % TFA), 50 µL Wash buffer 2 (10 % ACN, 0.2% TFA), and tips transferred to a fresh vial. Bound phosphopeptides were eluted by addition of 30 µL Elution buffer 1 (1 % NH_4_OH), directly centrifuged and the eluate transferred to a fresh vial to be directly acidified with 30 µL 10 % formic acid to minimize deamidation and loss of phosphorylation. The elution step was repeated with 30 µL Elution buffer 2 (5 % NH_4_OH, 25 % ACN). The same protocol was repeated a second time with a different loading buffer - Loading Buffer 2 containing lactic acid (20 % lactic acid, 70 % ACN, 5 % TFA) - to maximize the phosphorylated peptide yield.

### Mass spectrometric analysis

Peptides were resuspended in 3 % ACN, 0.1 % formic acid and approximately 500 ng loaded onto a 50 cm Acclaim PepMap column using an EASY-nLC 1200 system coupled to a QExactive HF mass spectrometer. For interactome analysis, peptides were resolved across a 218 min active gradient at a 150 nL/min flow rate. Mass spectrometric analysis was done in data-independent mode using variable windows to achieve similarly complex fragment mass spectra across the entire range of 300-1650 m/z using 24 windows with 1 m/z overlaps (Bruderer *et al*., 2017). Full mass spectra were recorded in the Orbitrap at a resolution of 120K in a mass range of 300-1650 m/z with a maximum injection time (max. IT) of 60 ms and an AGC target of 3e6. Precursor ions were isolated in the quadrupole and fragmented with stepped HCD at 28 ± 3 % NCE, before fragment mass spectra were scanned in the Orbitrap at a resolution of 30K, an AGC target of 1e6 while max. IT was set to auto. For phosphopeptide samples, peptides were resolved across a 150 min gradient, and DIA measurement with 20 variable windows was performed in a mass range of 350-1600 m/z. All other settings were similar as described for the interactome fraction.

### Data analysis

Raw data were analyzed with Spectronaut v18.3.230830.50606 in directDIA mode. For analysis of the interactome fraction, default BGS factory settings were used except minimum peptide length was set to 6. For analysis of the two phosphoenriched fractions, BGS factory settings were used with min. 6 aa, variable modifications set to include phosphorylation at S, T, Y, deamidation at N, Q, as well as glutamine to pyro-glutamate modification. The Normalization Type Filter was set to include phosphorylated residues. PTM workflow was enabled and the localization probability filter was set to 0.75. Only proteins that were confidently identified with a Q-value ≤ 0.01 were retained. Quantification results were exported as pivot reports for statistical analysis with Perseus (Tyanova *et al*., 2016). For the interactome fraction, results were log2 transformed and filtered for hits which were identified and quantified in at least 3 of the 4 biological replicates (relaxed filter) or in all 4 replicates, as well as identified with >1 precursor ion (strict filter). Then, missing values were imputed using a tail-based imputation. To determine differences between more than two conditions, multiple-sample t-tests were performed with S0=0 and FDR ≤ 0.01, only significant hits kept, values z-score normalized and the results visualized in a hierarchical cluster analysis using the Euclidean distance. For further statistical analysis of inhibitor-related differences, two-sample t-tests were performed with S0 = 0.1 and FDR ≤ 0.05 for the pairwise comparisons of LRRK2 with MLi-2 against LRRK2, LRRK1 with IN04 against LRRK1, LRRK1 or LRRK2 against Control and LRRK2 versus LRRK1. Then, results were filtered for significant hits in the LRRK1 and LRRK2 conditions, z-score normalized and the results displayed by hierarchical cluster analysis. Phosphoenrichment fraction results were filtered to keep only phosphosite identified in >50% of the replicates, imputed and normalized using the phosR package (Kim *et al*., 2021). Further statistical analysis was done in Perseus where multiple-sample tests (S0=0, FDR ≤ 0.05), and two-sample tests (S0=0.1, FDR ≤ 0.05) were performed. Results were each filtered for significant hits and z-score normalized. Multiple-sample test results were visualized as heatmap while two-sample test results were displayed as volcano plots.

### Protein Network Analysis

For further interactome analysis, cluster derived from the hierarchical cluster analysis were put into cytoscape (v3.10.3) and networks obtained using the STRING: protein query application. Displayed edges have a confidence score of at least 0.8 and interactions were analyzed by MCL clustering with clustermaker2 (v2.3.4.; (Utriainen and Morris, 2023)). Nodes were analysed for functional enrichment of Gene Ontology cellular compartment annotations in the cluster against the human genome. Nodes were colored continuously after the specific strength of the enrichment (as determined by the Posthoc test). Node size was assigned to confidence of the hit identification as determined by the C-score of Spectronaut. Volcano and PTMvsProtein fold change plots were produced with python (version 3.9.13) using the pandas (version 1.4.4, https://pandas.pydata.org/) and the plotly express (plotly version 5.9.0, https://plotly.com/python/plotly-express/) packages.

## References

Adams, J.A. (2001) ‘Kinetic and catalytic mechanisms of protein kinases’, Chemical Reviews, 101(8), pp. 2271–2290. Available at: 10.1021/CR000230W/ASSET/IMAGES/LARGE/CR000230WF00011.JPEG.

Bae, E.J. and Lee, S.J. (2020) ‘The LRRK2-RAB axis in regulation of vesicle trafficking and α-synuclein propagation’, Biochimica et Biophysica Acta (BBA) - Molecular Basis of Disease, 1866(3), p. 165632. Available at: 10.1016/J.BBADIS.2019.165632.

Barrett, J.C. et al. (2008) ‘Genome-wide association defines more than 30 distinct susceptibility loci for Crohn’s disease’, Nature Genetics, 40(8), pp. 955–962. Available at: 10.1038/ng.175.

Beilina, A. et al. (2014) ‘Unbiased screen for interactors of leucine-rich repeat kinase 2 supports a common pathway for sporadic and familial Parkinson disease’, Proceedings of the National Academy of Sciences of the United States of America, 111(7), pp. 2626–2631. Available at: 10.1073/pnas.1318306111.

Beilina, A. et al. (2020) ‘The Parkinson’s Disease Protein LRRK2 Interacts with the GARP Complex to Promote Retrograde Transport to the trans-Golgi Network’, Cell Reports, 31(5), p. 107614. Available at: 10.1016/J.CELREP.2020.107614.

Berginski, M.E. et al. (2021) ‘The Dark Kinase Knowledgebase: an online compendium of knowledge and experimental results of understudied kinases’, Nucleic Acids Research, 49(D1), pp. D529–D535. Available at: 10.1093/NAR/GKAA853.

Biskup, S. et al. (2006) ‘Localization of LRRK2 to membranous and vesicular structures in mammalian brain’, Annals of Neurology, 60(5), pp. 557–569. Available at: 10.1002/ana.21019.

Biskup, S. et al. (2007) ‘Dynamic and redundant regulation of LRRK2 and LRRK1 expression’, BMC Neuroscience, 8(1), pp. 1–11. Available at: 10.1186/1471-2202-8-102/FIGURES/4.

Blanca-Ramírez, M. et al. (2017) ‘GTP binding regulates cellular localization of Parkinson’s disease-associated LRRK2’, Human Molecular Genetics, 26(14), pp. 2747–2767. Available at: 10.1093/HMG/DDX161.

Bruderer, R. et al. (2017) ‘Optimization of Experimental Parameters in Data-Independent Mass Spectrometry Significantly Increases Depth and Reproducibility of Results’, Molecular & Cellular Proteomics, 16(12), p. 2296. Available at: 10.1074/MCP.RA117.000314.

Cho, K.F. et al. (2020) ‘Proximity labeling in mammalian cells with TurboID and split-TurboID’, Nature Protocols [Preprint]. Available at: 10.1038/s41596-020-0399-0.

Civiero, L. et al. (2012) ‘Biochemical Characterization of Highly Purified Leucine-Rich Repeat Kinases 1 and 2 Demonstrates Formation of Homodimers’, PLoS ONE, 7(8). Available at: 10.1371/JOURNAL.PONE.0043472.

Cui, L. et al. (2022) ‘Vesicle trafficking and vesicle fusion: mechanisms, biological functions, and their implications for potential disease therapy’, Molecular Biomedicine. Springer. Available at: 10.1186/s43556-022-00090-3.

Deng, X. et al. (2011) ‘Characterization of a selective inhibitor of the Parkinson’s disease kinase LRRK2’, Nature Chemical Biology *2011 7:4*, 7(4), pp. 203–205. Available at: 10.1038/nchembio.538.

Deniston, C.K. et al. (2020) ‘Structure of LRRK2 in Parkinson’s disease and model for microtubule interaction’, Nature, 588(7837), pp. 344–349. Available at: 10.1038/S41586-020-2673-2.

Dzamko, N. et al. (2010) ‘Inhibition of LRRK2 kinase activity leads to dephosphorylation of Ser(910)/Ser(935), disruption of 14-3-3 binding and altered cytoplasmic localization’, The Biochemical Journal, 430(3), pp. 405–413. Available at: 10.1042/BJ20100784.

Erb, M.L. and Moore, D.J. (2020) ‘LRRK2 and the Endolysosomal System in Parkinson’s Disease’, Journal of Parkinson’s Disease, 10, pp. 1271–1291. Available at: 10.3233/JPD-202138.

Fell, M.J. et al. (2015) ‘MLi-2, a Potent, Selective, and Centrally Active Compound for Exploring the Therapeutic Potential and Safety of LRRK2 Kinase Inhibition’, J Pharmacol Exp Ther, 355, pp. 397–409. Available at: 10.1124/jpet.115.227587.

Feng, D.D., Cai, W. and Chen, X. (2015) ‘The associations between Parkinson’s disease and cancer: the plot thickens’, Translational Neurodegeneration, 4(1). Available at: 10.1186/S40035-015-0043-Z.

di Fonzo, A. et al. (2005) ‘A frequent LRRK2 gene mutation associated with autosomal dominant Parkinson’s disease’, Lancet (London, England), 365(9457), pp. 412–415. Available at: 10.1016/S0140-6736(05)17829-5.

Gialluisi, A. et al. (2021) ‘Identification of sixteen novel candidate genes for late onset Parkinson’s disease’, Molecular Neurodegeneration, 16(1), pp. 1–18. Available at: 10.1186/S13024-021-00455-2/FIGURES/6.

Gilks, W.P. et al. (2005) ‘A common LRRK2 mutation in idiopathic Parkinson’s disease’, Lancet (London, England), 365(9457), pp. 415–416. Available at: 10.1016/S0140-6736(05)17830-1.

Godena, V.K. et al. (2014) ‘Increasing microtubule acetylation rescues axonal transport and locomotor deficits caused by LRRK2 Roc-COR domain mutations’, Nature Communications, 5. Available at: 10.1038/ncomms6245.

Haft, C.R. et al. (2000) ‘Human orthologs of yeast vacuolar protein sorting proteins Vps26, 29, and 35: assembly into multimeric complexes’, Molecular biology of the cell, 11(12), pp. 4105– 4116. Available at: 10.1091/MBC.11.12.4105.

Han, K.A. et al. (2017) ‘Leucine-rich repeat kinase 2 exacerbates neuronal cytotoxicity through phosphorylation of histone deacetylase 3 and histone deacetylation’, Human Molecular Genetics, 26(1), pp. 1–18. Available at: 10.1093/hmg/ddw363.

Hanafusa, H. et al. (2011) ‘Leucine-rich repeat kinase LRRK1 regulates endosomal trafficking of the EGF receptor’, Nature Communications, 2(1), p. 158. Available at: 10.1038/NCOMMS1161.

Hanafusa, H. et al. (2015) ‘PLK1-dependent activation of LRRK1 regulates spindle orientation by phosphorylating CDK5RAP2’, Nature cell biology, 17(8), pp. 1024–1035. Available at: 10.1038/NCB3204.

Hanley, S.E. and Cooper, K.F. (2021) ‘Sorting nexins in protein homeostasis’, Cells. MDPI, pp. 1–26. Available at: 10.3390/cells10010017.

Härtlova, A. et al. (2018) ‘LRRK2 is a negative regulator of Mycobacterium tuberculosis phagosome maturation in macrophages’, The EMBO Journal, 37(12). Available at: 10.15252/EMBJ.201798694.

Herzog, D. et al. (2022) ‘Chemical Proteomics of the Tumor Suppressor Fhit Covalently Bound to the Cofactor Ap3A Elucidates Its Inhibitory Action on Translation’, Journal of the American Chemical Society, 144(19), pp. 8613–8623. Available at: 10.1021/JACS.2C00815/ASSET/IMAGES/LARGE/JA2C00815_0008.JPEG.

Ho, D.H. et al. (2015) ‘Leucine-Rich Repeat Kinase 2 (LRRK2) phosphorylates p53 and induces p21(WAF1/CIP1) expression’, Molecular brain, 8(1). Available at: 10.1186/S13041-015-0145-7.

Hu, J. et al. (2023) ‘Small-molecule LRRK2 inhibitors for PD therapy: Current achievements and future perspectives’, European Journal of Medicinal Chemistry, 256, p. 115475. Available at: 10.1016/j.ejmech.2023.115475.

Hui, K.Y. et al. (2018) ‘Functional variants in the LRRK2 gene confer shared effects on risk for Crohn’s disease and Parkinson’s disease’, Science Translational Medicine, 10(423). Available at: 10.1126/SCITRANSLMED.AAI7795.

Ishikawa, K. et al. (2012) ‘EGFR-dependent phosphorylation of leucine-rich repeat kinase LRRK1 is important for proper endosomal trafficking of EGFR’, Molecular Biology of the Cell, 23(7), pp. 1294–1306. Available at: 10.1091/MBC.E11-09-0780/ASSET/IMAGES/LARGE/1294FIG8.JPEG.

Kalogeropulou, A.F. et al. (2018) ‘P62/SQSTM1 is a novel leucine-rich repeat kinase 2 (LRRK2) substrate that enhances neuronal toxicity’, The Biochemical journal, 475(7), pp. 1271–1293. Available at: 10.1042/BCJ20170699.

Kalogeropulou, A.F. et al. (2022) ‘Impact of 100 LRRK2 variants linked to Parkinson’s Disease on kinase activity and microtubule binding’, Biochemical Journal, 479(17), pp. 1759–1783. Available at: 10.1042/BCJ20220161/935911/bcj-2022-0161.pdf.

Kedashiro, S. et al. (2015) ‘LRRK1-phosphorylated CLIP-170 regulates EGFR trafficking by recruiting p150Glued to microtubule plus ends’, Journal of Cell Science, 128(2), pp. 385–396. Available at: 10.1242/JCS.161547/VIDEO-7.

Kim, H.J. et al. (2021) ‘PhosR enables processing and functional analysis of phosphoproteomic data’, Cell Reports, 34(8), p. 108771. Available at: 10.1016/J.CELREP.2021.108771.

Kim, T. et al. (2019) ‘HDAC inhibition by valproic acid induces neuroprotection and improvement of PD-like behaviors in LRRK2 R1441G transgenic mice’, Experimental Neurobiology, 28(4), pp. 504–515. Available at: 10.5607/en.2019.28.4.504.

Kobayashi, T. et al. (2011) ‘Centriolar Kinesin Kif24 Interacts with CP110 to Remodel Microtubules and Regulate Ciliogenesis’, Cell, 145(6), pp. 914–925. Available at: 10.1016/J.CELL.2011.04.028.

Kuwahara, T. et al. (2016) ‘LRRK2 and RAB7L1 coordinately regulate axonal morphology and lysosome integrity in diverse cellular contexts’, Scientific Reports, 6. Available at: 10.1038/SREP29945.

Law, B.M.H. et al. (2014) ‘A Direct Interaction between Leucine-rich Repeat Kinase 2 and Specific β-Tubulin Isoforms Regulates Tubulin Acetylation’, Journal of Biological Chemistry, 289(2), pp. 895–908. Available at: 10.1074/JBC.M113.507913.

Lebovitz, C. et al. (2021) ‘Loss of Parkinson’s susceptibility gene LRRK2 promotes carcinogen-induced lung tumorigenesis’, Scientific Reports, 11(1). Available at: 10.1038/S41598-021-81639-0.

Lesage, S. et al. (2007) ‘LRRK2 exon 41 mutations in sporadic Parkinson disease in Europeans’, Archives of neurology, 64(3), pp. 425–430. Available at: 10.1001/ARCHNEUR.64.3.425.

MacLeod, D.A. et al. (2013) ‘RAB7L1 Interacts with LRRK2 to Modify Intraneuronal Protein Sorting and Parkinson’s Disease Risk’, Neuron, 77(3), pp. 425–439. Available at: 10.1016/J.NEURON.2012.11.033.

Malik, A.U. et al. (2021) ‘Deciphering the LRRK code: LRRK1 and LRRK2 phosphorylate distinct Rab proteins and are regulated by diverse mechanisms’, Biochemical Journal, 478(3), pp. 553–578. Available at: 10.1042/BCJ20200937.

Manschwetus, J.T. et al. (2020) ‘Binding of the Human 14-3-3 Isoforms to Distinct Sites in the Leucine-Rich Repeat Kinase 2’, Frontiers in neuroscience, 14. Available at: 10.3389/FNINS.2020.00302.

Marín, I. (2006) ‘The Parkinson disease gene LRRK2: evolutionary and structural insights’, Molecular Biology and Evolution, 23(12), pp. 2423–2433. Available at: 10.1093/MOLBEV/MSL114.

Meixner, A. et al. (2011) ‘A QUICK Screen for Lrrk2 Interaction Partners-Leucine-rich Repeat Kinase 2 is Involved in Actin Cytoskeleton Dynamics’, Molecular and Cellular Proteomics, 10(1). Available at: 10.1074/mcp.M110.001172.

Myasnikov, A. et al. (2021) ‘Structural analysis of the full-length human LRRK2’, Cell, 184(13), pp. 3519–3527.e10. Available at: 10.1016/J.CELL.2021.05.004.

Nagy, Z. et al. (2010) ‘The metazoan ATAC and SAGA coactivator HAT complexes regulate different sets of inducible target genes’, Cellular and molecular life sciences : CMLS, 67(4), pp. 611–628. Available at: 10.1007/S00018-009-0199-8.

Nalls, M.A., et al. (2014) ‘NeuroGenetics Research Consortium (NGRC) 19, Hussman Institute of Human Genomics (HIHG) 19, The Ashkenazi Jewish Dataset Investigator 19, Cohorts for Health and Aging Research in Genetic Epidemiology (CHARGE) 19, North American Brain Expression Consortium (NABEC) 19, United Kingdom Brain Expression Consortium (UKBEC) 19, Greek Parkinson’s Disease Consortium 19’, Nature Publishing Group, 46(9), p. 36. Available at: 10.1038/ng.3043.

Nichols, R.J. et al. (2010) ‘14-3-3 binding to LRRK2 is disrupted by multiple Parkinson’s disease-associated mutations and regulates cytoplasmic localization’, Biochemical Journal, 430(3), pp. 393–404. Available at: 10.1042/BJ20100483.

Obsilova, V. and Obsil, T. (2022) ‘Structural insights into the functional roles of 14-3-3 proteins’, Frontiers in Molecular Biosciences. Frontiers Media S.A. Available at: 10.3389/fmolb.2022.1016071.

Orpinell, M. et al. (2010) ‘The ATAC acetyl transferase complex controls mitotic progression by targeting non-histone substrates’, The EMBO journal, 29(14), pp. 2381–2394. Available at: 10.1038/EMBOJ.2010.125.

Paisán-Ruíz, C. et al. (2004) ‘Cloning of the Gene Containing Mutations that Cause PARK8-Linked Parkinson’s Disease’, Neuron, 44(4), pp. 595–600. Available at: 10.1016/J.NEURON.2004.10.023.

Perez-Riverol, Y. et al. (2022) ‘The PRIDE database resources in 2022: a hub for mass spectrometry-based proteomics evidences’, Nucleic Acids Research, 50(D1), p. D543. Available at: 10.1093/NAR/GKAB1038.

Rappsilber, J., Mann, M. and Ishihama, Y. (2007) ‘Protocol for micro-purification, enrichment, pre-fractionation and storage of peptides for proteomics using StageTips’, Nature Protocols, 2(8), pp. 1896–1906. Available at: 10.1038/NPROT.2007.261;KWRD=LIFE+SCIENCES.

Reimer, J.M. et al. (2023) ‘Structure of LRRK1 and mechanisms of autoinhibition and activation’, Nature Structural & Molecular Biology *2023 30:11*, 30(11), pp. 1735–1745. Available at: 10.1038/s41594-023-01109-1.

Reyniers, L. et al. (2014) ‘Differential protein-protein interactions of LRRK1 and LRRK2 indicate roles in distinct cellular signaling pathways’, Journal of Neurochemistry, 131(2), p. 239. Available at: 10.1111/JNC.12798.

Rojas, R. et al. (2008) ‘Regulation of retromer recruitment to endosomes by sequential action of Rab5 and Rab7’, Journal of Cell Biology, 183(3), pp. 513–526. Available at: 10.1083/jcb.200804048.

Saha, M. et al. (2012) ‘RSK phosphorylates SOS1 creating 14-3-3 docking sites and negatively regulating MAPK activation’, The Biochemical journal, 447(1), p. 159. Available at: 10.1042/BJ20120938.

Sakane, A. et al. (2021) ‘JRAB/MICAL-L2 undergoes liquid-liquid phase separation to form tubular recycling endosomes’, Communications biology, 4(1). Available at: 10.1038/S42003-021-02080-7.

Salašová, A. et al. (2017) ‘A proteomic analysis of LRRK2 binding partners reveals interactions with multiple signaling components of the WNT/PCP pathway’, Molecular Neurodegeneration, 12(1). Available at: 10.1186/s13024-017-0193-9.

Sanger, A. et al. (2019) ‘Adaptor protein complexes and disease at a glance’, J Cell Sci., 132(20). Available at: 10.1242/jcs.222992.

Saunders-Pullman, R. et al. (2010) ‘LRRK2 G2019S Mutations are associated with an increased cancer risk in Parkinson Disease’, Movement Disorders, 25(15), p. 2536. Available at: 10.1002/MDS.23314.

Schindler, C. et al. (2015) ‘EARP is a multisubunit tethering complex involved in endocytic recycling’, Nature Cell Biology *2014 17:5*, 17(5), pp. 639–650. Available at: 10.1038/ncb3129.

Schmidt, S.H. et al. (2019) ‘The dynamic switch mechanism that leads to activation of LRRK2 is embedded in the DFGψ motif in the kinase domain’, Proceedings of the National Academy of Sciences of the United States of America, 116(30), pp. 14979–14988. Available at: 10.1073/PNAS.1900289116/-/DCSUPPLEMENTAL.

Schneider, C.A., Rasband, W.S. and Eliceiri, K.W. (2012) ‘NIH Image to ImageJ: 25 years of image analysis’, Nature Methods *2012 9:7*, 9(7), pp. 671–675. Available at: 10.1038/nmeth.2089.

Seaman, M.N.J. (2012) ‘The retromer complex-endosomal protein recycling and beyond’, Journal of Cell Science, pp. 4693–4702. Available at: 10.1242/jcs.103440.

Shu, L. et al. (2019) ‘A comprehensive analysis of population differences in LRRK2 variant distribution in Parkinson’s disease’, Frontiers in Aging Neuroscience, 11(JAN). Available at: 10.3389/FNAGI.2019.00013/FULL.

Si, M. et al. (2019) ‘A small molecular inhibitor of LRRK1 identified by homology modeling and virtual screening suppresses osteoclast function, but not osteoclast differentiation, in vitro’, Aging (Albany NY), 11(10), p. 3250. Available at: 10.18632/AGING.101977.

Sommese, R.F. and Sivaramakrishnan, S. (2016) ‘Substrate Affinity Differentially Influences Protein Kinase C Regulation and Inhibitor Potency’, The Journal of Biological Chemistry, 291(42), p. 21963. Available at: 10.1074/JBC.M116.737601.

Steger, M. et al. (2016) ‘Phosphoproteomics reveals that Parkinson’s disease kinase LRRK2 regulates a subset of Rab GTPases’, eLife, 5(January). Available at: 10.7554/ELIFE.12813.001.

Steger, M. et al. (2017) ‘Systematic proteomic analysis of LRRK2-mediated rab GTPase phosphorylation establishes a connection to ciliogenesis’, eLife, 6. Available at: 10.7554/ELIFE.31012.

Stein, M.P. et al. (2003) ‘Human VPS34 and p150 are Rab7 interacting partners’, Traffic, 4(11), pp. 754–771. Available at: 10.1034/J.1600-0854.2003.00133.X.

Stormo, A.E.D. et al. (2022) ‘The E3 ligase TRIM1 ubiquitinates LRRK2 and controls its localization, degradation, and toxicity’, The Journal of Cell Biology, 221(4). Available at: 10.1083/JCB.202010065.

Tasegian, A. et al. (2021) ‘Impact of Type II LRRK2 inhibitors on signaling and mitophagy’, Biochemical Journal, 478(19), pp. 3555–3573. Available at: 10.1042/BCJ20210375.

Taylor, S.S. et al. (2020) ‘Kinase Domain Is a Dynamic Hub for Driving LRRK2 Allostery’, Frontiers in Molecular Neuroscience, 13, p. 538219. Available at: 10.3389/FNMOL.2020.538219/BIBTEX.

Terai, T. et al. (2006) ‘JRAB/MICAL-L2 Is a Junctional Rab13-binding Protein Mediating the Endocytic Recycling of Occludin’, Molecular Biology of the Cell, 17(5), p. 2465. Available at: 10.1091/MBC.E05-09-0826.

Tomkins, J.E. et al. (2018) ‘Comparative Protein Interaction Network Analysis Identifies Shared and Distinct Functions for the Human ROCO Proteins’, Proteomics, 18(10). Available at: 10.1002/PMIC.201700444.

Tong, Y. et al. (2012) ‘Loss of leucine-rich repeat kinase 2 causes age-dependent bi-phasic alterations of the autophagy pathway’, Molecular Neurodegeneration, 7(1). Available at: 10.1186/1750-1326-7-2.

Tyanova, S., et al. (2016) ‘The Perseus computational platform for comprehensive analysis of (prote)omics data’, Nature Methods. Nature Publishing Group, pp. 731–740. Available at: 10.1038/nmeth.3901.

Ungar, D. et al. (2002) ‘Characterization of a mammalian Golgi-localized protein complex, COG, that is required for normal Golgi morphology and function’, Journal of Cell Biology, 157(3), pp. 405–415. Available at: 10.1083/jcb.200202016.

Ungar, D. et al. (2006) ‘Retrograde transport on the COG railway’, Trends in Cell Biology, pp. 113–120. Available at: 10.1016/j.tcb.2005.12.004.

Usmani, A., Shavarebi, F. and Hiniker, A. (2021) ‘The Cell Biology of LRRK2 in Parkinson’s Disease’, Molecular and Cellular Biology, 41(5), pp. e00660–20. Available at: 10.1128/MCB.00660-20.

Utriainen, M. and Morris, J.H. (2023) ‘clusterMaker2: a major update to clusterMaker, a multi-algorithm clustering app for Cytoscape’, BMC Bioinformatics, 24(1), pp. 1–28. Available at: 10.1186/S12859-023-05225-Z/FIGURES/10.

Wang, D. et al. (2015) ‘Association of the LRRK2 genetic polymorphisms with leprosy in Han Chinese from Southwest China’, Genes and Immunity, 16(2), pp. 112–119. Available at: 10.1038/GENE.2014.72.

Wang, Z. et al. (2018) ‘Meta-analysis of human gene expression in response to Mycobacterium tuberculosis infection reveals potential therapeutic targets’, BMC systems biology, 12(1). Available at: 10.1186/S12918-017-0524-Z.

Wauters, L., Versées, W. and Kortholt, A. (2019) ‘Roco Proteins: GTPases with a Baroque Structure and Mechanism’, International Journal of Molecular Sciences, 20(1). Available at: 10.3390/IJMS20010147.

Wei, Y. et al. (2023) ‘The function of Golgi apparatus in LRRK2-associated Parkinson’s disease’, Frontiers in Molecular Neuroscience, 16, p. 1097633. Available at: 10.3389/FNMOL.2023.1097633/BIBTEX.

West, A.B. et al. (2005) ‘Parkinson’s disease-associated mutations in leucine-rich repeat kinase 2 augment kinase activity’, Proceedings of the National Academy of Sciences of the United States of America, 102(46), pp. 16842–16847. Available at: 10.1073/PNAS.0507360102/SUPPL_FILE/07360FIG8.JPG.

Westerlund, M. et al. (2008) ‘Developmental regulation of leucine-rich repeat kinase 1 and 2 expression in the brain and other rodent and human organs: Implications for Parkinson’s disease’, Neuroscience, 152(2), pp. 429–436. Available at: 10.1016/j.neuroscience.2007.10.062.

Willett, R., Ungar, D. and Lupashin, V. (2013) ‘The Golgi puppet master: COG complex at center stage of membrane trafficking interactions’, Histochemistry and Cell Biology, 140(3), pp. 271–283. Available at: 10.1007/S00418-013-1117-6/FIGURES/4.

Xing, W.R. et al. (2017) ‘Role and mechanism of action of leucine-rich repeat kinase 1 in bone’, Bone research, 5, p. 17003. Available at: 10.1038/BONERES.2017.3.

Yaffe, M.B. et al. (1997) ‘The Structural Basis for 14-3-3:Phosphopeptide Binding Specificity’, Cell, 91(7), pp. 961–971. Available at: 10.1016/S0092-8674(00)80487-0.

Yamamura, R. et al. (2008) ‘The Interaction of JRAB/MICAL-L2 with Rab8 and Rab13 Coordinates the Assembly of Tight Junctions and Adherens Junctions’, Molecular Biology of the Cell, 19, pp. 971–983.

Yang, Z. et al. (2022) ‘Multifaceted Roles of Retromer in EGFR Trafficking and Signaling Activation’, Cells, 11(21). Available at: 10.3390/cells11213358.

Zeng, C. et al. (2016) ‘Leucine-rich repeat kinase-1 regulates osteoclast function by modulating RAC1/Cdc42 Small GTPase phosphorylation and activation’, American journal of physiology. Endocrinology and metabolism, 311(4), pp. E772–E780. Available at: 10.1152/AJPENDO.00189.2016.

Zhang, F.-R. et al. (2009) ‘Genomewide association study of leprosy’, The New England Journal of Medicine, 361(27), pp. 2609–2618. Available at: 10.1056/NEJMOA0903753.

Zhang, T. et al. (2022) ‘Interrogating Kinase−Substrate Relationships with Proximity Labeling and Phosphorylation Enrichment’, Journal of Proteome Research, 21, pp. 494–506. Available at: 10.1021/acs.jproteome.1c00865.

Zimprich, A. et al. (2004) ‘Mutations in LRRK2 cause autosomal-dominant parkinsonism with pleomorphic pathology’, Neuron, 44(4), pp. 601–607. Available at: 10.1016/j.neuron.2004.11.005.

